# A parental transcriptional response to microsporidia infection induces inherited immunity in offspring

**DOI:** 10.1101/2020.10.11.335117

**Authors:** Alexandra R. Willis, Winnie Zhao, Ronesh Sukhdeo, Lina Wadi, Hala Tamim El Jarkass, Julie M. Claycomb, Aaron W. Reinke

## Abstract

Inherited immunity is an emerging field and describes how the transfer of immunity from parents to offspring can promote progeny survival in the face of infection. The mechanisms of how inherited immunity is induced are mostly unknown. The intracellular parasite *Nematocida parisii* is a natural microsporidian pathogen of *Caenorhabditis elegan*s. Here, we show that *N. parisii*-infected worms produce primed offspring that are resistant to microsporidia infection. We find that immunity is induced in a dose dependent manner and lasts for a single generation. Intergenerational immunity prevents host cell invasion by *N. parisii* and also enhances survival to the bacterial pathogen *Pseudomonas aeruginosa*. Further, we show that inherited immunity is triggered by the host transcriptional response to infection, which can also be induced through maternal somatic depletion of negative regulators PALS-22 and the retinoblastoma protein ortholog LIN-35. We show that other biotic and abiotic stresses, such as viral infection and cadmium exposure, that induce a similar transcriptional response to microsporidia can also induce immunity in progeny. Our results demonstrate that distinct stimuli can induce inherited immunity to provide resistance against multiple classes of pathogens. These results show that activation of an innate immune response can provide protection against pathogens not only within a generation, but also in the next generation.

## Introduction

Animals have evolved diverse immune mechanisms to limit the negative impact of pathogens and parasites on host fitness. While immunological memory is typically considered a hallmark of the antibody-mediated adaptive immune system, memory of pathogen exposure has now been documented in animals lacking adaptive immunity. Although, invertebrates only posses innate immunity, at least 20 species, including insects, crustaceans, and molluscs, have now been shown to transfer protective immunity to their progeny following infection^1^. Although this epigenetically inherited immunity can protect offspring against a variety of bacterial, fungal and viral pathogens, it is largely unclear how immunity is induced. Several reports have described the deposition of bacterial cell-wall fragments in offspring following parental infection, as well as immune genes being upregulated in both parents and their progeny^2,3^. Immune priming can be specific, whereby immunity is only active against the same strain of bacteria that the parents were infected with^4,5^. Conversely, it may be broad; for example, mealworm beetles primed with either fungi or a Gram-positive or Gram-negative bacteria induce immunity against Gram-positive pathogens^6^. Although the effectors that provide immunity in the progeny are mostly unknown, antimicrobial peptides are often upregulated in offspring^1,2,6^.

Studies in the genetically tractable nematode *C. elegans* have enabled fundamental immune advances and shed light on many epigenetically inherited and stress-induced phenotypes^7–9^. As such, *C. elegans* has recently become a powerful model for the study of inherited innate immunity^10^. In this host, antiviral immunity, mediated by the small RNA mediated silencing of viral transcripts, was shown to last for at least three generations and be dependent on RNA interference (RNAi) pathways^11^. Although there are conflicting reports about whether the natural Orsay virus can induce heritable immunity in *C. elegans*, injection with vesicular stomatitis virus was able to protect progeny against reinfection^12–14^. Several studies have shown that learned bacterial avoidance can be transferred to progeny. In one case, heritable avoidance to *Pseudomonas aeruginosa* bacteria was dependent on RNAi pathways and a bacterial RNA used by *C. elegans* to inhibit host gene expression^15–17^. Parental exposure to pathogenic bacteria can also protect offspring by increasing the likelihood of progeny adopting a stress-resistant dauer phenotype^18^. Finally, resistance can be induced by upregulating immune genes in offspring. For example, intergenerational immunity to pathogenic *Pseudomonas vranovensis* is dependent on the induced expression of cysteine synthases^19,20^.

Microsporidia are a large phylum of fungal-related parasites that infect most animal species^21^. These pathogens can be lethal to immunocompromised humans, and can cause huge economic losses by infecting agriculturally important animals such as honey bees, silk worms and shrimp^22^. *Nematocida parisii* is a natural microsporidia parasite that commonly infects *C. elegans* in the wild. Infection of *C. elegans* by *N. parisii* begins when microsporidia spores are ingested^23^. Spores inside the intestinal lumen then fire a unique polar-tube structure which is used to transfer the cellular contents of the spore (sporoplasm), including the parasite’s genetic material into the host intestinal cell^24^. The pathogen then replicates intracellularly to form meronts, and spreads from cell to cell by fusing the cells and forming syncytia^25^. These meronts ultimately differentiate into mature spores which exit infected cells non-lytically, and can result in the shedding of up to 200,000 spores during infection^23,26^. Microsporidia are pathogenic to *C. elegans*, resulting in reduced fecundity and ultimately death^25,27^.

Several mechanisms of innate immunity against microsporidia have been described in invertebrates. Microsporidia infection typically induces a strong host transcriptional response, which often includes upregulation of a suite of different antimicrobial peptides^28^. Though antimicrobial peptides are commonly upregulated in other invertebrates, and *C. elegans* possess many different families of these proteins, so far only C-type lectins have been shown to be upregulated upon *N. parisii* infection^29,30^. Instead, *C. elegans* possesses a novel stress/immune pathway called the ‘intracellular pathogen response’ (IPR) that is induced upon infection by both microsporidia and Orsay virus^29,31^. The IPR includes upregulation of ubiquitin adapter proteins and is thought is to be involved in clearing intracellular parasites^29,32,33^.

In this study, we reveal robust resistance to microsporidia in the offspring of *N. parisii*-infected *C. elegans*. We show that inherited immunity dramatically reduces host-cell invasion by *N. parisii* and also confers resistance to the bacterial pathogen *P. aeruginosa*. We find that immunity is conferred in a dose-dependent manner, lasts throughout development, and is maintained for a single generation. Using tissue-specific depletion or expression of negative regulators of the IPR (LIN-35 and PALS-22), we demonstrate that immunity in progeny is dependent on a parental transcriptional response, and that this immunity is transferred from maternal somatic tissues to the progeny. Remarkably, the host transcriptional response and thus inherited immunity, can be induced by both biotic and abiotic stresses which mimic the microsporidial response. Together, these results provide insight into how inherited immune responses can be induced to provide immunity against pathogenic infection.

## 2. Results

### 2.1. Parental infection by *N. parisii* confers immunity to *C. elegans* progeny in a dose-dependent manner

Infection of *C. elegans* with the natural microsporidian pathogen *N. parisii* delays development and impacts worm fertility^24,34^. To quantify the effects of infection, we exposed early larval stage (L1) animals to varying doses of *N. parisii* spores. At 72 hours post infection (hpi), animals were fixed and stained with the chitin-binding dye DY96 (Figures S1A and S1B). DY96 stains both microsporidia spores and worm embryos, allowing us to determine both parasite burden and host reproductive development. Worms infected with higher doses of spores exhibited greater parasite burden and a smaller body size at 72 hpi (Figures S1C and S1D). Higher infection doses also correlated with a reduction in the percentage of gravid adults (animals carrying embryos), as well as a reduction in the number of embryos per animal (Figures S1E and S1F). In agreement with these findings, infection of transgenic animals expressing a fluorescent protein in the germline revealed germline deformities in worms exposed to high doses of spores (Figure S1G).

To determine the effects of parental infection on offspring, we infected parental (P0) generations at the L1 stage with a ‘very low’, ‘low’ or ‘moderate’ dose of *N. parisii* that resulted in a ∼10-50% reduction in the number of gravid adults (Figure S1E). At 72 hpi, infected adults were treated with a sodium hypochlorite solution to release the F1 embryos from adults and destroy microsporidia spores. As *N. parisii* infection is not vertically transmitted, resultant larvae are not infected with microsporidia (Figure S2)^27^. F1 generations of synchronized L1s were then exposed to a high dose of spores. Remarkably, imaging revealed a quantifiable reduction in the parasite burden of primed worms from infected parents, as compared to naïve worms from uninfected parents (Figures 1A and 1B). Immunity in F1 progeny was dose-dependent; parents experiencing a higher infection burden gave rise to offspring that were more resistant to *N. parisii*. In agreement with our data for parasite burden, we saw that primed worms under infection conditions were larger and produced significantly more embryos than their naïve counterparts (Figures 1C-1E). Primed worms showed no fitness advantage under non-infection conditions and were indeed smaller and contained fewer embryos than worms from uninfected parents (Figures S3A and S3B). Together, these data reveal a robust and dose-dependent immunity phenotype in the offspring of *N. parisii*-infected worms.

**Figure 1.**
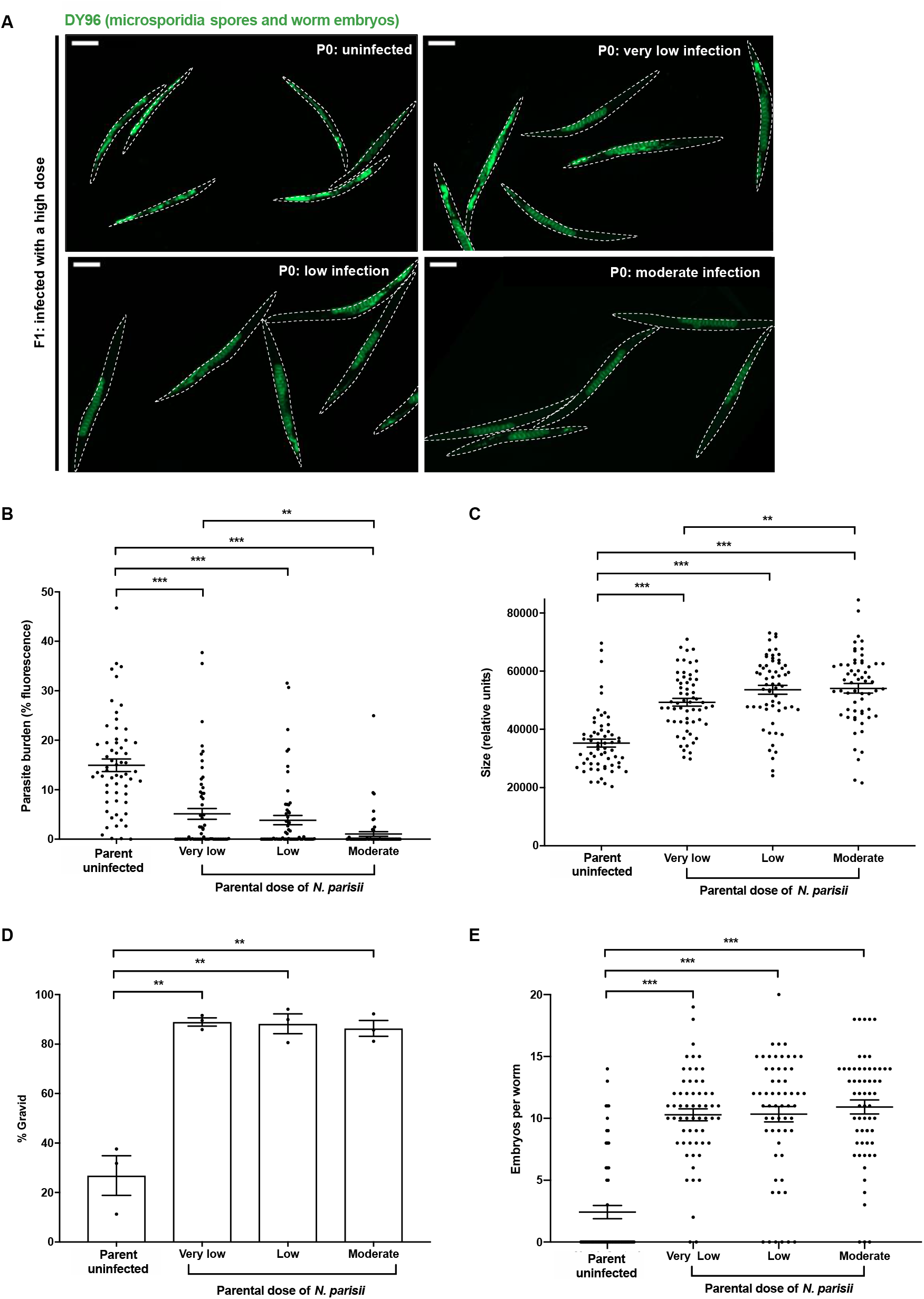
Parental infection by *N. parisii* confers immunity to the progeny of *C. elegans*. P0 populations of N2 *C. elegans* were uninfected or exposed to varying concentrations of *N. parisii* spores at the L1 stage (doses defined in methods). At 72 hpi, animals were treated with sodium hypochlorite solution to release F1 embryos. F1 L1 larvae were then exposed to a high dose of *N. parisii*. At 72 hpi F1 animals were fixed and stained with DY96 to visualize both *N. parisii* spores and worm embryos. (A) Representative images of F1 populations stained with DY96. Scale bars, 200 μm. (B) Images of DY96 stained F1 worms were analysed and fluorescence from *N. parisii* spores thresholded to determine parasite burdens of individual worms (% of body filled with spores). Each circle represents a measurement from a single worm. Mean ± SEM (horizontal bars) is shown. Data pooled from 3 independent experiments using n = 20 worms per condition per experiment. (C) Images of F1 worms were analysed, and the area of individual worms calculated. Each circle represents a measurement from a single worm. Mean ± SEM (horizontal bars) is shown. Data pooled from 3 independent experiments using n = 20 worms per condition per experiment. (D) Images of DY96 stained F1 worms were analysed and worms possessing 1 or more embryos were considered gravid. Mean ± SEM (horizontal bars) is shown. Data pooled from 3 independent experiments using n = 98-351 worms per condition per experiment. (E) Images of DY96 stained F1 worms were analysed and embryos per worm quantified. Each circle represents a count from a single worm. Mean ± SEM (horizontal bars) is shown. Data pooled from 3 independent experiments using n = 20 worms per condition per experiment. The p-values were determined by unpaired two-tailed Student’s t-test. (B-E) Significance with Bonferroni correction was defined as p < 0.0166. **, p < 0.0033; ***, p < 0.00033.

### 2.2 Inherited immunity prevents microsporidia invasion events by restricting spores in the intestine

To resist microsporidia infection, animals may (i) prevent invasion of intestinal cells or (ii) destroy the invaded pathogen^24,33^. To determine by which mechanism inherited immunity confers resistance to *N. parisii*, we first performed invasion assays. Naïve or primed L1 worms were exposed to a high dose of spores and fixed 30 minutes post infection (mpi) or 3 hpi. To visualize invasion events in these animals, we performed fluorescence *in situ* hybridization (FISH) to detect *N. parisii* 18S ribosomal RNA in host intestinal cells, thus indicating the presence of intracellular sporoplasms. Our analysis revealed significantly fewer invasion events in primed animals at both time points, including an over 98% reduction in infectious events at 30 mpi (Figures 2A, 2B and S4A). To determine if spores were present in the intestinal lumen, animals were also stained with DY96. Strikingly, we also observed far fewer spores in the guts of primed animals (Figures 2C, 2D and S4B).

**Figure 2.**
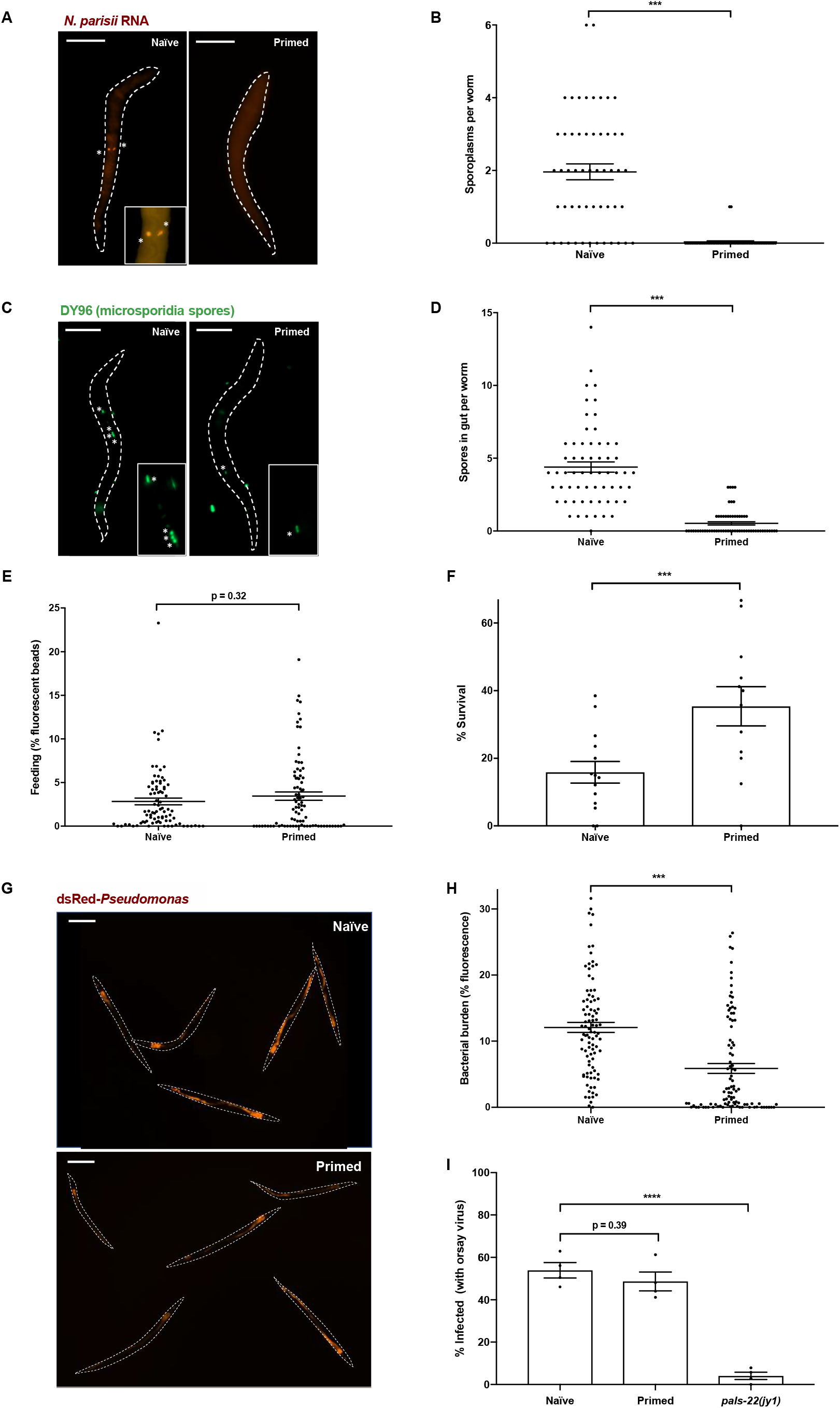
Inherited immunity prevents microsporidia invasion and *P. aeruginosa* colonization, but not viral infection. (A-D) P0 populations of N2 *C. elegans* were either not infected or infected with a low dose of *N. parisii* spores at the L1 stage (doses defined in methods). At 72 hpi, animals were treated with sodium hypochlorite solution to release F1 embryos. Naïve and primed F1 larvae were exposed to a maximal dose of *N. parisii* at the L1 stage. At 30 mpi, animals were fixed and stained with DY96 to detect *N. parisii* spores (green) as well as a FISH probe to detect *N. parisii* 18S RNA (red). (A) Representative images of worms stained with FISH probe to detect invaded sporoplasms, marked by asterisks. Scale bars, 25 μm. (B) The number of sporoplasms per animal was quantified by microscopy. Each circle represents a count from a single worm. Mean ± SEM (horizontal bars) is shown. Data pooled from 3 independent experiments using n = 16-20 worms per condition per experiment. (C) Representative images of worms stained with DY96 to detect spores, marked by asterisks, in the intestinal lumen. Scale bars, 25 μm. (D) The number of spores per animal was quantified by microscopy. Each circle represents a count from a single worm. Mean ± SEM (horizontal bars) is shown. Data pooled from 3 independent experiments using n = 20-21 worms per condition per experiment. (E) Naïve and primed F1 larvae were fed fluorescent beads at the L1 stage. After 30 min, animals were fixed and imaged. Fluorescence from beads was thresholded to determine the amount of beads eaten by each worm (% of body filled with beads). Each circle represents a measurement from a single worm. Mean ± SEM (horizontal bars) is shown. Data pooled from 3 independent experiments using n = 24-30 worms per condition per experiment. (F) Naïve and *N. parisii*-primed F1 L1 larvae were maintained on slow-killing plates with wild-type *P. aeruginosa* and survival monitored. Animal survival at an 84 hpi end point is shown. Mean ± SEM (horizontal bars) is shown. Data pooled from 4 independent experiments each comprising 2-4 technical replicates, using n = 13-37 worms per condition per experiment. (G-H) Naïve and *N. parisii*-primed F1 L1 larvae were maintained on slow-killing plates with dsRed-*P. aeruginosa* and fixed at 48 hpi. (G) Representative images of worm populations grown on dsRed-*P. aeruginosa*. Scale bars, 200 μm. (H) Images of worms were analysed and fluorescence thresholded to determine bacterial burdens of individual worms (% of body filled with *P. aeruginosa*). Each circle represents a measurement from a single worm. Mean ± SEM (horizontal bars) is shown. Data pooled from 3 independent experiments using n = 21-47 worms per condition per experiment. (I) Naïve and primed F1, and *pals-22* mutant L1 larvae were infected with Orsay virus. At 16 hpi, worms were fixed and stained with FISH probe to detect Orsay virus RNA. Worms with one or more cells stained by FISH were counted as infected. Mean ± SEM (horizontal bars) is shown. Data pooled from 4 independent experiments with n > 52 worms per condition per experiment. The p-values were determined by unpaired two-tailed Student’s t-test. (B-F, H) Significance was defined as p < 0.05; **, p < 0.01; ***, p < 0.001. (I) Significance with Bonferroni correction was defined as *, p < 0.025; ****, p < 0.0001.

To test whether a reduction of spores in our primed worms was a result of reduced feeding, we allowed naïve and primed L1 stage worms to feed on fluorescent beads. After 30 min or 3 h, we fixed the populations and quantified fluorescence in individual animals. We failed to detect any reduction in feeding in primed worms at either time point (Figures 2E and S4C). At the 3 h time point, primed animals actually fed significantly more than naïve worms, indicating that microsporidia invasion results in reduced feeding in *C. elegans* (Figure S4C).

To test whether enhanced clearance of intracellular infection might also contribute to microsporidia resistance in primed animals, we performed pulse-chase experiments^24^. Here, naïve and primed animals were maintained in the presence of high concentrations of *N. parisii* spores for 3 h, before washing thoroughly to remove any microsporidia spores not inside the animals. The population was then split; half was immediately fixed to represent the initial infection, and the other half maintained in the absence of spores until a 24 h end point when they were fixed to assess clearance. Detection of sporoplasms using 18S RNA FISH and subsequent quantifications showed no evidence of intracellular pathogen clearance in the primed animals (Figure S4D). These data suggest that limiting invasion is the principal way by which inherited immunity provides protection against microsporidia.

To provide insight into the mechanisms underlying protection in primed animals, we next tested the specificity of the immune response. For this, we assayed resistance to the extracellular Gram-negative bacterial pathogen *Pseudomonas aeruginosa* (Strain PA14) using a well-established ‘slow killing’ protocol^7^. Here, naïve or *N. parisii*-primed animals were plated on wild-type PA14 at the L1 stage and survival assayed at 84 hpi. Remarkably, primed worms were significantly less susceptible to *P. aeruginosa* infection than naïve animals (Figure 2F). Slow killing by *P. aeruginosa* occurs as a result of bacterial accumulation in the gut^35^. To visualize bacterial burden in our naïve and primed worms, we performed infection assays using dsRed-expressing PA14. At 48 hpi, we fixed and quantified fluorescence in our naïve and primed populations. In agreement with the data from survival assays, the bacterial burden in microsporidia-primed animals was significantly reduced (Figures 2G, 2H and S4E). Together, these results suggest that a host intestinal factor may be protecting primed worms from infection by destroying microsporidia spores and other pathogenic microbes within the intestinal lumen.

To test whether inherited immunity could protect against all classes of pathogen, we tested the response of *N. parisii*-primed animals to the intracellular intestinal pathogen Orsay virus. Surprisingly, FISH staining revealed that primed worms were similarly susceptible to Orsay virus as their naïve counterparts (Figure 2I). This result is particularly notable given the significant overlap between the transcriptional responses induced by Orsay virus and *N. parisii*, with both activating IPR genes^36^. PALS-22 is a negative regulator of the IPR and a loss-of-function mutation in *pals-22* results in animals with a constitutively activated IPR response that are resistant to viral and microsporidia infection, but not to *P. aeruginosa* (Figures 2I and S4F)^32^. Mutants deficient for *pals-22* limit microsporidia infection by preventing invasion as well as clearing invaded parasites (Figure S4F). Thus, IPR-mediated immunity is distinct in several ways from the immunity found in primed animals.

### 2.3 Inherited immunity to *N. parisii* lasts a single generation and persists throughout development

We next sought to understand the kinetics of the inherited immune response to microsporidia infection. To determine whether immunity could be transmitted over multiple generations, we tested resistance to microsporidia in both the F1 and F2 progeny of *N. parisii*-infected worms. We observed resistance to microsporidia only in the F1 population, indicating that this response lasts for a single generation (Figures 3A and 3B, S5A and S5B). To test the longevity of the inherited immune response within the F1 generation, naïve and primed worms were infected with *N. parisii* either at the L1 stage, 24 h later at the L2/L3 stage, or 48 h later at the L4 stage. After 30 mpi, worms were fixed, stained with a FISH probe to detect *N. parisii*, and the number of sporoplasms per individual quantified. While immune-primed worms continued to show some resistance to infection at the L4 stage, resistance was strongest at the L1 stage of infection (Figure 3C).

**Figure 3.**
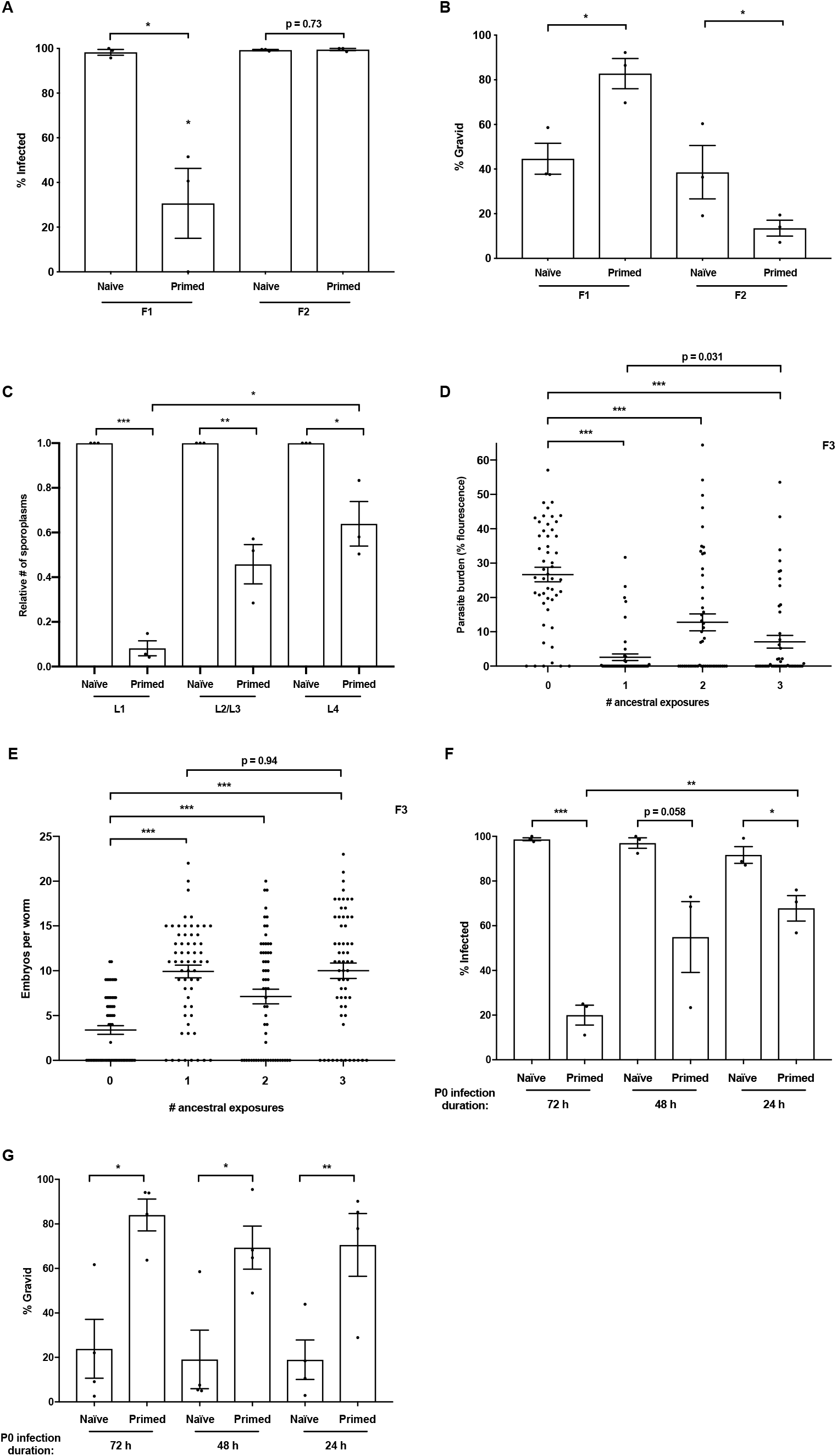
Inherited immunity in *N. parisii* primed *C. elegans* lasts a single generation and is strongest in early larval stages. (A-B) P0 populations of N2 *C. elegans* were either not infected or infected with a low dose of *N. parisii* spores at the L1 stage (doses defined in methods). At 72 hpi, animals were treated with sodium hypochlorite solution to release F1 embryos. F1 embryo populations were then split and either tested for immunity, or maintained under non-infection conditions for the collection and subsequent testing of F2 embryos. Both naïve and primed F1 and F2 larvae were exposed to a high dose of *N. parisii* at the L1 stage. At 72 hpi, F1 and F2 animals were fixed and stained with DY96 to visualize *N. parisii* spores and embryos. (A) Individual DY96 stained worms were imaged to determine infection status. Mean ± SEM (horizontal bars) is shown. Data pooled from 3 independent experiments using n = 16-20 worms per condition per experiment. (B) Images of DY96 stained worms were analysed and worms possessing 1 or more embryos were considered gravid. Mean ± SEM (horizontal bars) is shown. Data pooled from 3 independent experiments using n = 16-20 worms per condition per experiment. (C) Naïve and primed F1 larvae were obtained as above and challenged with a maximal dose of *N. parisii* spores at either the L1, L2/L3 or L4 stage. Animals were fixed at 30 mpi and stained with a FISH probe to detect *N. parisii* 18S RNA. The number of sporoplasms per animal was quantified by microscopy. Shown is the average number of sporoplasms in primed worms relative to the naïve control. Mean ± SEM (horizontal bars) is shown. Data pooled from 3 independent experiments using n = 12-20 worms per condition per experiment. (D-E) Briefly, N2 *C. elegans* (P0, F1, F2) were infected at the L1 stage with *N. parisii* for one, two or three successive generations. Each infection period lasted 72 h before treating with sodium hypochlorite solution to obtain the next generation of embryos (see schematic Figure S5C). F3 larvae were exposed to a high dose of *N. parisii* at the L1 stage. At 72 hpi, F3 animals were fixed and stained with DY96 to visualize *N. parisii* spores and embryos. (D) Images of DY96 stained F3 worms were analysed and fluorescence from *N. parisii* spores thresholded to determine parasite burdens of individual worms (% of body filled with spores). Each circle represents a measurement from a single worm. Mean ± SEM (horizontal bars) is shown. Data pooled from 2 independent experiments using n = 25 worms per condition per experiment. (E) Images of DY96 stained F3 worms were analysed and embryos per worm quantified. Each circle represents a count from a single worm. Mean ± SEM (horizontal bars) is shown. Data pooled from 2 independent experiments using n = 30 worms per condition per experiment. (F-G) P0 populations of N2 *C. elegans* were either not infected or infected with a low dose of *N. parisii* spores at the L1 stage (for 72 h), L2/L3 stage (for 48 h) or L4 stage (for 24 h). At 72 h post L1, animals were treated with sodium hypochlorite solution to release F1 embryos. Both naïve and primed F1 larvae were exposed to a high dose of *N. parisii* at the L1 stage. At 72 hpi, animals were fixed and stained with DY96 to visualize *N. parisii* spores and embryos. (F) Individual DY96 stained worms were imaged to determine infection status. Mean ± SEM (horizontal bars) is shown. Data pooled from 3 independent experiments using n = 100-197 worms per condition per experiment. (G) Images of DY96 stained worms were analysed and worms possessing 1 or more embryos were considered gravid. Mean ± SEM (horizontal bars) is shown. Data pooled from 4 independent experiments using n = 100-197 worms per condition per experiment. The p-values were determined by unpaired two-tailed Student’s t-test. (A-B, G) Significance was defined as: *, p < 0.05; **, p < 0.01; ***, p < 0.001. (C, F) Significance with Bonferroni correction was defined as: *, p < 0.025; **, p < 0.005; ***, p < 0.0005. (D-E). Significance with Bonferroni correction was defined as: *, p < 0.016.

We next tested whether a greater ancestral history of infection might enhance immunity phenotypes and potentially enable the immune response to transmit over multiple generations. Animals were infected for one, two, or three sequential generations with *N. parisii* (P0s, F1s and F2s). Susceptibility to microsporidia was assessed in F3 animals, as well as in F4 animals following a single generation of rest without infection (Figures S5C-E). We found that an increased history of ancestral infections did not enhance immunity phenotypes among the F3 populations (Figures 3D and 3E). Furthermore, resistance was seen exclusively in the F3 animals and no resistance was observed in the F4 generations (Figures S5F and S5G).

We next wanted to determine for how long a parent must be infected with *N. parisii* in order to transmit immunity to progeny. For this, parental generations were infected as previously at the L1 stage for 72 h of total infection, or rested on their typical *E. coli* food source and subsequently infected as L2/L3s for 48 h total infection, or as L4s for 24 h total infection. Infection assays on resultant F1 progenies showed that parental worms infected only briefly as L4s were still able to confer immunity phenotypes to offspring, though to a lesser extent than animals from parents infected for longer periods (Figures 3F and 3G). As *N. parisii* takes over 48 hours to sporulate^25^, these results indicate that the early stages of microsporidian infection alone (i.e. invasion and replication) are sufficient to induce immunity in progeny.

Together, these data show that inherited immunity protects the F1 progeny from infected parents throughout development and supports a model in which the immediate parental environment is the most important factor in determining the immune competency of offspring.

### 2.4 The parental transcriptional response to infection triggers inherited immunity in offspring

*N. parisii* is an intestinal parasite that does not come into direct contact with the *C. elegans* germline, and how information is transferred from the soma to the germline is not known (Figure S6A). To explore this, we tested whether mutants defective in small-RNA inheritance and histone modification were able to transmit immunity phenotypes to offspring. We found that the offspring of *N. parisii* infected mutants still became protected against infection (Figure S7A-D)^19,37^. In addition, we found that the master immune regulator PMK-1 was not required for the induction of immunity, consistent with this pathway not being involved in immunity to *N. parisii* (Figure S7C-D)^27^. These data indicate that the small RNA, histone modification, and PMK-1 pathways are not required for transmission of immunity.

To determine whether infection itself, or merely exposure to spores, is required for the transmission of inherited immunity from parent to progeny, we used heat-killed spores. Parent populations were either uninfected, exposed to live *N. parisii*, or exposed to heat-killed spores. Unlike live spores, heat-killed spores fail to induce the IPR response, as demonstrated using transcriptional reporters for key IPR genes (Figures S6B and S6C) ^36^. Consistent with a requirement for infection and induction of the IPR to initiate inherited immunity, the offspring of parents exposed to heat-killed spores showed no enhanced protection against *N. parisii* infection (Figures 4A and 4B).

**Figure 4.**
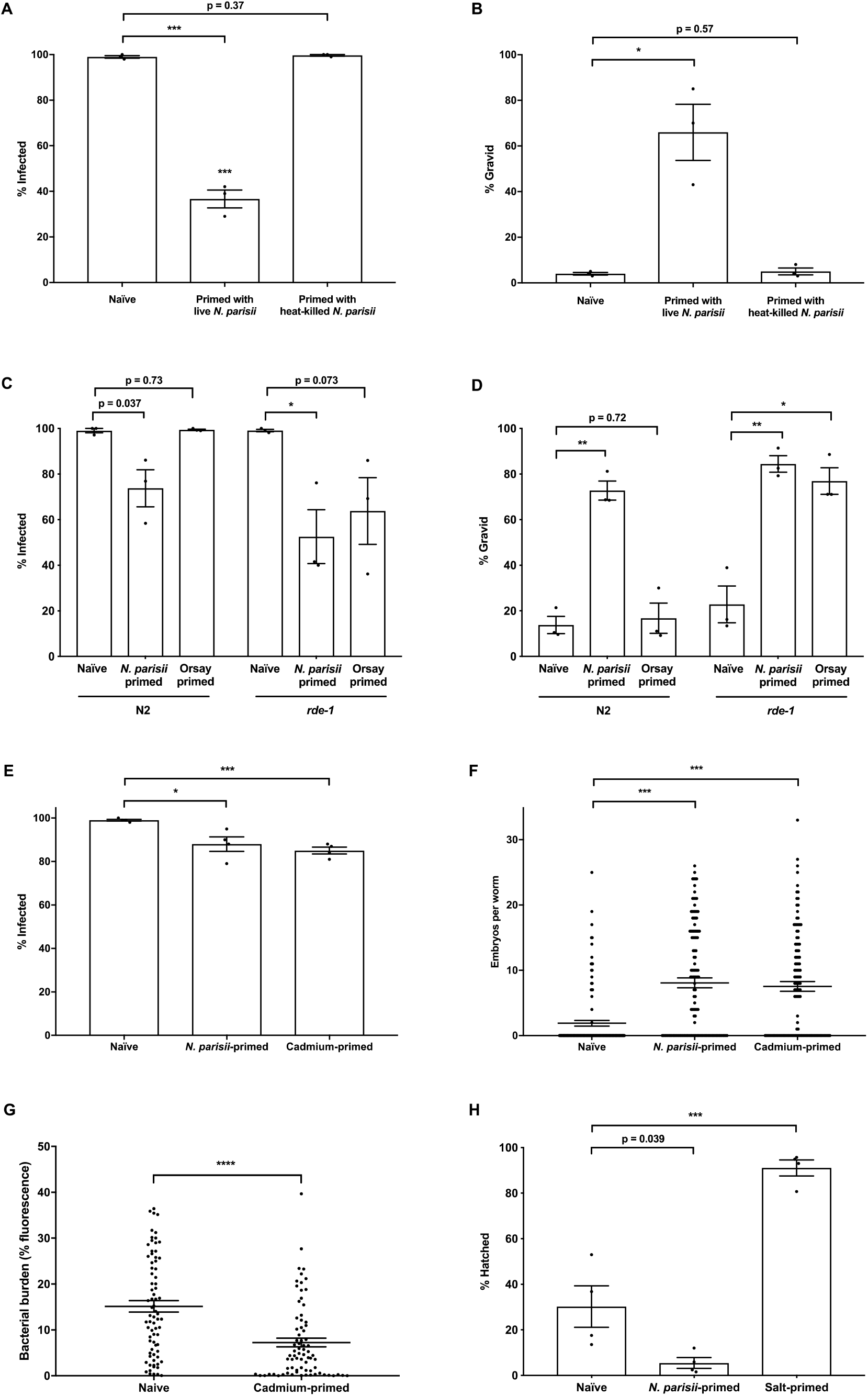
The parental transcriptional response to *N. parisii* triggers inherited immunity. (A-B) P0 populations of N2 *C. elegans* were either not infected or exposed to a low dose of heat-killed or live *N. parisii* spores at the L1 stage (doses defined in methods). At 72 hpi, animals were treated with sodium hypochlorite solution to release F1 embryos. F1 larvae were exposed to a high dose of *N. parisii* at the L1 stage. At 72 hpi, animals were fixed and stained with DY96 to visualize both *N. parisii* spores and worm embryos. (A) Individual DY96 stained worms were imaged to determine infection status. Mean ± SEM (horizontal bars) is shown. Data pooled from 3 independent experiments using n =100 worms per condition per experiment. (B) Images of DY96 stained F1 worms were analysed and worms possessing 1 or more embryos were considered gravid. Mean ± SEM (horizontal bars) is shown. Data pooled from 3 independent experiments using n = 100 worms per condition per experiment. (C-D) P0 populations of N2 and *rde-1* mutants were not infected, infected with *N. parisii* (low dose), or infected with Orsay virus at the L1 stage. At 72 hpi, animals were treated with sodium hypochlorite solution to release F1 embryos. F1 larvae were exposed to a high dose of *N. parisii* at the L1 stage, then fixed and stained at 72 hpi as above. (C) DY96 stained worms were analyzed to determine infection status. Mean ± SEM (horizontal bars) is shown. Data pooled from 3 independent experiments using n = 100 worms per condition per experiment. (D) Images of DY96 stained F1 worms were analysed and worms possessing 1 or more embryos were considered gravid. Mean ± SEM (horizontal bars) is shown. Data pooled from 3 independent experiments using n = 100 worms per condition per experiment. (E-G) P0 populations of N2 *C. elegans* were either untreated, exposed to 50 mM cadmium from the L4 stage, or infected with a low dose of *N. parisii* spores at the L4 stage. After 24 h, animals were treated with sodium hypochlorite solution to release F1 embryos. (E-F) F1 larvae were exposed to a high dose of *N. parisii* at the L1 stage. At 72 hpi, animals were fixed and stained with DY96 to visualize both *N. parisii* spores and worm embryos. (E) Individual DY96 stained worms were imaged to determine infection status. Mean ± SEM (horizontal bars) is shown. Data pooled from 4 independent experiments using n = 100 worms per condition per experiment. (F) Images of DY96 stained worms were analysed and embryos per worm quantified. Each circle represents a count from a single worm. Mean ± SEM (horizontal bars) is shown. Data pooled from 4 independent experiments using n = 25-30 worms per condition per experiment. (G) Naïve and cadmium-primed F1 L1 larvae were maintained on slow-killing plates with dsRed-*P. aeruginosa* and fixed at 48 hpi. Images of worms were analysed and fluorescence thresholded to determine bacterial burdens of individual worms (% of body filled with bacteria). Each circle represents a measurement from a single worm. Mean ± SEM (horizontal bars) is shown. Data pooled from 3 independent experiments using n = 25 worms per condition per experiment. (H) P0 populations of N2 *C. elegans* were either untreated or infected with a low dose of *N. parisii* spores at the L1 stage on regular NGM (containing 50 mM salt), or maintained on 250 mM salt. After 72 h, animals were treated with sodium hypochlorite solution to release F1 embryos. F1 embryos were maintained on 420 mM salt. At 48 hpi, embryo hatching was assessed by light microscopy. Mean ± SEM (horizontal bars) is shown. Data pooled from 4 independent experiments using n = 100 worms per condition per experiment. The p-values were determined by unpaired two-tailed Student’s t-test. (A-F, H) Significance with Bonferroni correction was defined as: *, p < 0.025; **, p < 0.005; ***, p < 0.0005. (G) Significance was defined as p < 0.05. ***, p < 0.001.

We next tested whether other environmental conditions that induce a similar transcriptional response as microsporidia infection could also induce immunity in progeny. When analysing mRNA sequencing data, we confirmed a previously noted similarity to the Orsay virus response, and also noticed that the *C. elegans* response to *N. parisii* infection overlaps significantly with that of worms exposed to the heavy metal cadmium (Figure S6D).

To explore a role for the transcriptional response in inducing inherited immunity, we first performed *N. parisii* infection assays on the offspring of untreated parents, or parents exposed to either *N. parisii* or Orsay virus. We saw that the Orsay-primed F1 progeny of loss-of-function *rde-1* mutants, but not N2 worms, showed significantly reduced infection and improved fitness under microsporidia infection conditions, as compared to naïve controls (Figures 4C and D). This difference between the *rde-1* genotype and N2 can be attributed to the increased susceptibility of *rde-1* mutants to Orsay virus, allowing the P0s to have an enhanced viral response (Figure S6E)^38^.

We next assayed the susceptibility of cadmium-primed animals to *N. parisii* infection. Strikingly, resistance to microsporidia was observed in the offspring of parents exposed to cadmium (Figures 4E and 4F). Additionally, cadmium-primed animals exposed to *P. aeruginosa* had significantly lower bacterial burden than naïve animals, supporting of a role for the transcriptional response in transmitting immunity against both these classes of pathogens (Figure 4G). Conversely, both infection-primed and cadmium-primed animals were less fit and displayed reduced gravidity when faced with a high concentration of cadmium (Figure S6F). Similarly, we found that while the progeny of osmotically stressed parents were better able to cope with a high salt stress, the offspring of *N. parisii*-infected animals were less viable under this stress condition (Figure 4H). These data indicate that primed animals are not simply more resistant to any given stress, but specifically to pathogenic *N. parisii* and *P. aeruginosa*. Furthermore, these data support a role for the transcriptional response in transmitting inherited immunity to offspring, and highlight the complex relationships that underlie responses to abiotic and biotic stresses across generations.

### 2.5. Progeny of animals with an artificially-activated transcriptional response are resistant to infection

In addition to the previously characterized *pals-22* mutant, we also observed that mutants of *lin-35*, the *C. elegans* ortholog of retinoblastoma protein (RB), induce a similar transcriptional response to microsporidia infection (Figure S6D)^39,40^. These mutants share extensive similarity with the transcriptional response to *N. parisii*, with 242 shared genes upregulated at least two-fold in both *lin-35* and *pals-22* as well as *N. parisii* infected worms (Figure S8). To determine whether activating this host response in the absence of infection or environmental stimuli would induce immunity, we performed infection assays in *pals-22* and *lin-35* mutants. Both *pals-22* and *lin-35* mutants have defects resulting in fewer embryos, even when grown under normal conditions (Figure 5A). Thus, the number of embryos per worm in these mutants during infection conditions is similar to wild type. However, in agreement with a role for the IPR response in restricting infection, both *pals-22* and *lin-35* mutants had a significantly lower parasite burden than wild-type animals (Figure 5B).

**Figure 5.**
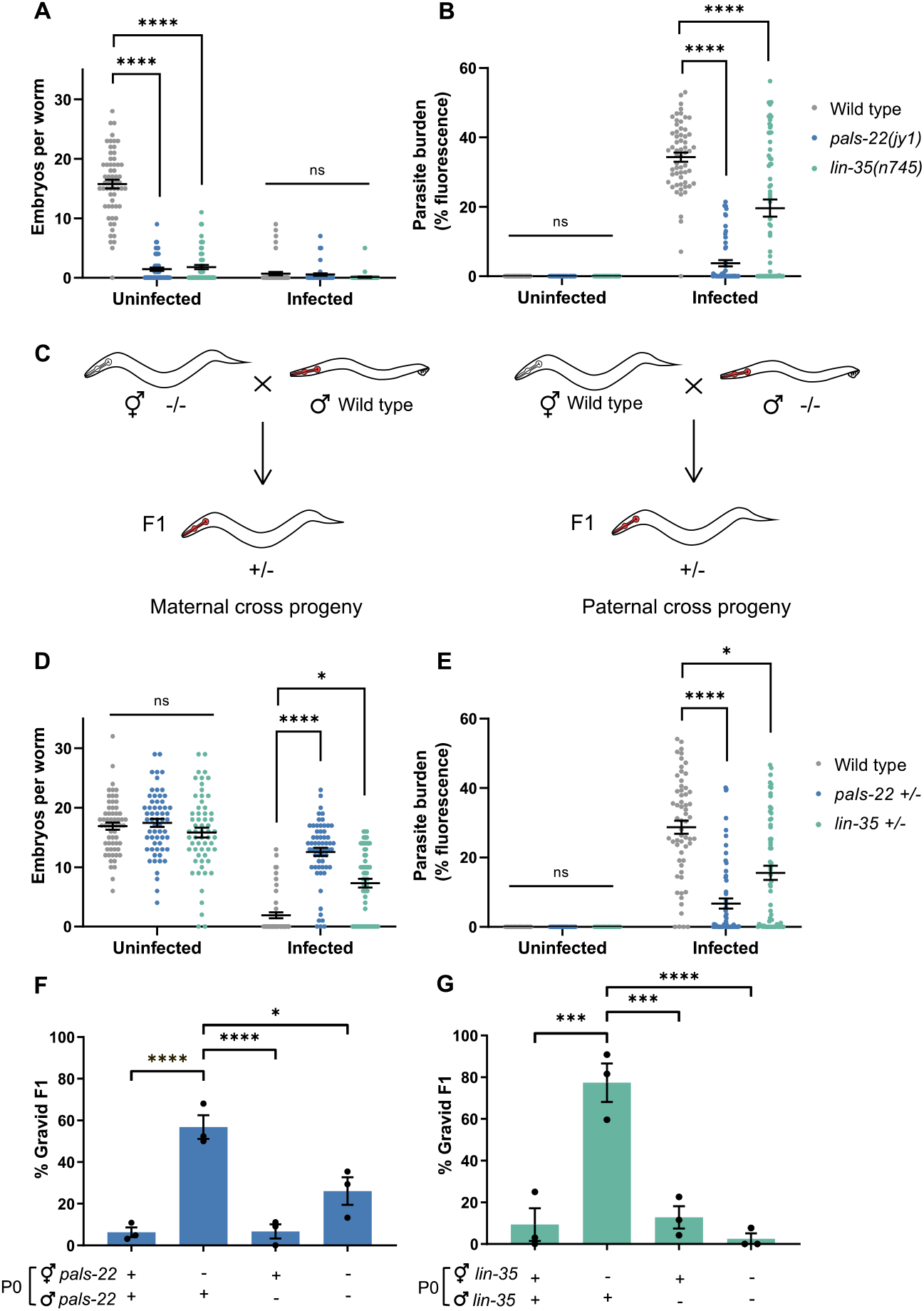
Mutants that phenocopy the transcriptional response to infection transfer immunity to offspring through the maternal germline. (A-B) Wild-type, *pals-22(jy1)*, and *lin-35(n745)* animals were not infected or infected with a high dose of *N. parisii* (infected) from the L1 stage. After 72 h, animals were fixed and stained with DY96 to visualize *N. parisii* spores and embryos. (A) Images of worms were analysed and embryos per worm quantified. Each circle represents a count from a single worm. Mean ± SEM (horizontal bars) is shown. Data pooled from 3 independent experiments using n = 20 worms per condition per experiment. (B) Images of worms were analysed and fluorescence from *N. parisii* spores thresholded to determine parasite burdens of individual worms (% of body filled with spores). Each circle represents a measurement from a single worm. Mean ± SEM (horizontal bars) is shown. Data pooled from 3 independent experiments using n = 20 worms per condition per experiment. (C) Schematic of mating to obtain maternal and paternal cross-progeny. *myo-2*p::mCherry used as a marker to distinguish cross-progeny from self-progeny. (D-E) P0 animals were allowed to mate for 24 h and then treated with sodium hypochlorite solution to release F1 embryos. F1 larvae were not infected or infected with a high dose of *N. parisii* at the L1 stage. At 72 hpi, animals were fixed and stained with DY96 to visualize *N. parisii* spores and embryos. (D) Images of worms were analysed and embryos per worm quantified. Each circle represents a count from a single worm. Mean ± SEM (horizontal bars) is shown. Data pooled from 3 independent experiments using n = 20 worms per condition per experiment. (E) Images of worms were analysed and fluorescence from *N. parisii* spores thresholded to determine parasite burdens of individual worms (% of body filled with spores). Each circle represents a measurement from a single worm. Mean ± SEM (horizontal bars) is shown. Data pooled from 3 independent experiments using n = 20 worms per condition per experiment. Only hermaphrodite maternal cross progeny are included in the quantifications. (F-G) P0 animals were allowed to mate for 24 h and then treated with sodium hypochlorite solution to release F1 embryos. F1 larvae were exposed to a high dose of *N. parisii* at the L1 stage. At 72 hpi, animals were fixed and stained with DY96 to visualize *N. parisii* embryos. Images of worms were analysed and worms possessing 1 or more embryos were considered gravid. Mean ± SEM (horizontal bars) is shown. Data pooled from 3 independent experiments using n = 13-67 worms per condition per experiment. The p-values were determined by Kruskal-Wallis test (non-parametric one-way ANOVA) (A-B, D-E) and by ordinary one-way ANOVA with post hoc (F-G). Significance was defined as: *, p < 0.05; **, p < 0.01; ***, p < 0.001, ****, p < 0.0001.

Next, to determine if transcriptional response activation in these mutants could induce immunity in progeny, we performed mating assays. We first examined the cross-progeny of *pals-22* and *lin-35* mutant hermaphrodites mated with wild-type males to determine if parents with an upregulated transcriptional response generated resistant F1s (Figure 5C). The heterozygous cross-progeny of both mutants produced a similar number of embryos in uninfected conditions as wild-type, as the developmental timing and brood size are recessive traits of *pals-22* and *lin-35* mutants. When infected with *N. parisii*, cross-progeny produced more embryos and have a lower pathogen load than wild-type animals (Figures 5D and 5E). On average, 57% of *pals-22* and 77% of *lin-35* maternal cross-progeny became gravid under infection conditions, compared to less than 10% of wild-type animals (Figures 5F and 5G). In contrast, paternal cross-progeny i.e. the offspring of wild-type hermaphrodites mated with males carrying either *pals-22* or *lin-35* mutations, do not exhibit improved reproductive fitness under infection conditions (Figures 5F and 5G). These results reveal that inherited immunity to microsporidia is maternally transferred and can be induced by transcriptional activation alone.

### 2.6. The signal for offspring resistance can originate in multiple somatic tissues

To identify the tissues required to induce the transcriptional response and thereby transmit inherited immunity, we used two methods. First, we infected the maternal cross-progeny of *pals-22* mutants carrying a transgene for wild-type *pals-22* under an endogenous promoter, or under a promoter specific for a single tissue type where *pals-22* is typically expressed^41^. While offspring of the *pals-22* endogenous rescue strain were no longer resistant to infection, neuronal and hypodermal-specific rescue strains still produced resistant progeny, but to a lesser extent than *pals-22* mutants (Figure 6A). This indicates that a signal in the neuronal and hypodermal tissues may contribute to the transmission of inherited immunity, but may not be crucial. The intestinal-specific rescue produced progeny that were less resistant than *pals-22* mutants, though this was not statistically significant, indicating that an intestinal signal could also contribute to transmission of immunity. The maternal cross-progeny of *lin-35* mutants with wild-type *lin-35* expressed under either an endogenous or intestine-specific promoter were no longer resistant to *N. parisii*, indicating that an intestinal signal was important for transmission of immunity in this case. Furthermore, addition of *lin-35* under the *pie-1* germline promoter was seen to partially reduce resistance (Figure 6B).

**Figure 6.**
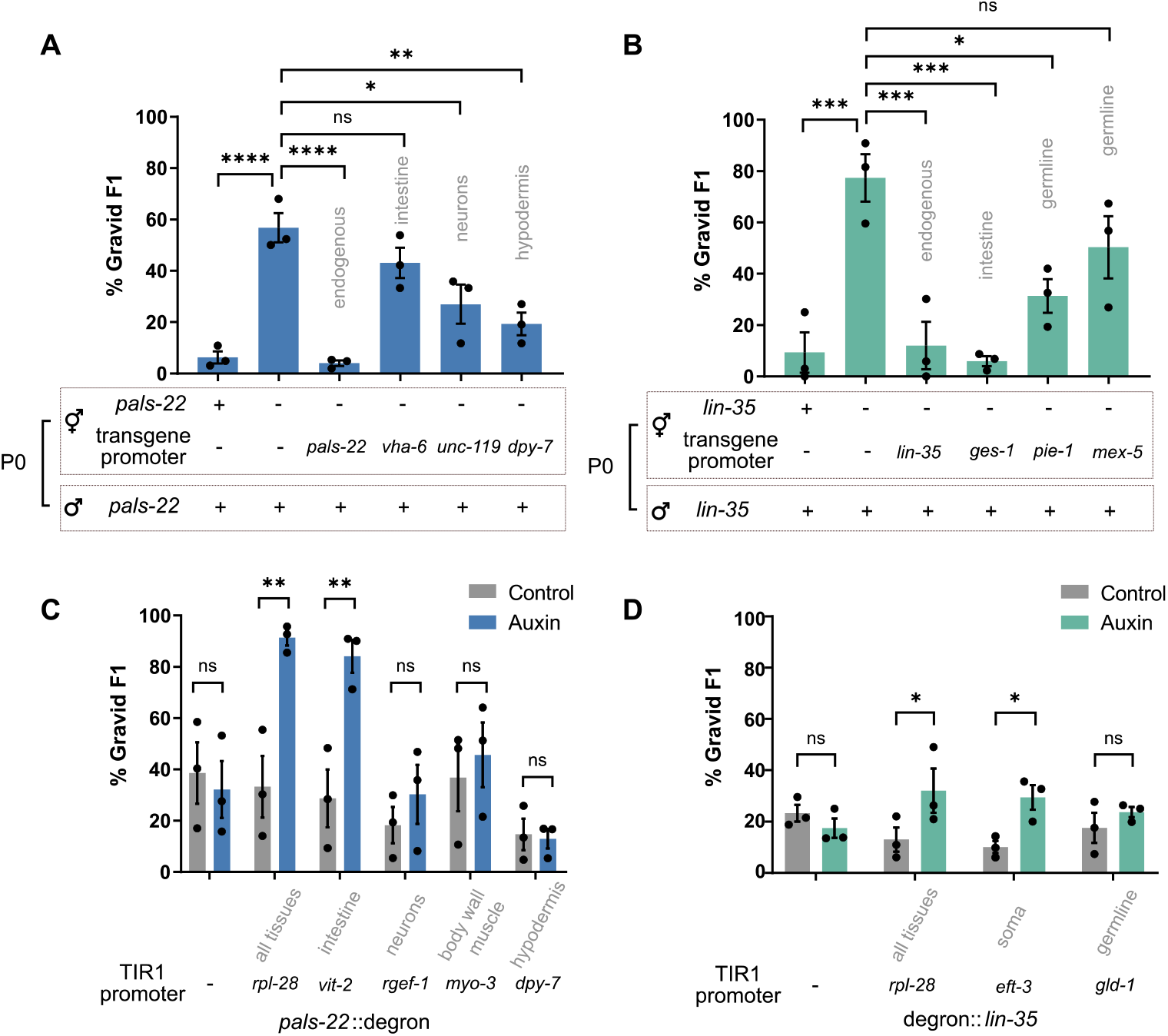
Induction of the IPR in somatic tissues induces inherited immunity. (A-B) P0 animals were allowed to mate for 24 h and then treated with sodium hypochlorite solution to release F1 embryos. F1 larvae were exposed to a high dose of *N. parisii* at the L1 stage. At 72 hpi, animals were fixed and stained with DY96 to visualize embryos. The percentage of gravid F1 was quantified for the maternal cross progeny of (A) pals-22 mutants carrying a wild-type pals-22 transgene and (B) lin-35 mutants carrying a wild-type lin-35 transgene. Mean ± SEM (horizontal bars) is shown. Data pooled from 3 independent experiments using n = 17-78 worms per condition per experiment. Tissues expressing the rescue transgenes are indicated on the graph. (C-D) P0 animals were grown on either control plates or plates containing 200uM auxin from embryos to mediate degradation of PALS-22 or LIN-35. At 72 h animals were treated with sodium hypochlorite solution to release F1 embryos. F1 larvae were exposed to a high dose of *N. parisii* at the L1 stage. At 72 hpi, animals were fixed and stained with DY96 to visualize embryos. The percentage of gravid F1 was quantified for (C) degron-tagged pals-22 strains and (D) degron-tagged lin-35 strains. Mean ± SEM (horizontal bars) is shown. Data pooled from 3 independent experiments using n > 100 worms per condition per experiment. Tissues expressing the rescue transgenes are indicated on the graph. The p-values were determined by ordinary one-way ANOVA with post hoc (A-D). Significance was defined as: *, p < 0.05; **, p < 0.01; ***, p < 0.001; ****, p < 0.0001.

In a second approach, we used the auxin inducible degradation (AID) system^42^ to degrade either PALS-22 or LIN-35 in a tissue-specific manner in the parental generation only, and assessed F1 progenies for resistance to microsporidia (Figure S9). Targeted degradation of PALS-22, and thus signal induction, in the adult intestine alone was sufficient to enhance immunity in progeny, while degradation in any other single tissue was unable to provide protection (Figure 6C). Additionally, degradation of LIN-35 in somatic tissues only, was sufficient to confer resistance to offspring (Figure 6D). Taken together, these results suggest that induced expression of the IPR either in the adult intestine only, or in multiple somatic tissues is sufficient to transmit inherited immunity to offspring.

## 3. Discussion

Inherited immunity is a nascent and rapidly growing field of research, with important consequences for our understanding of health and evolution. Multiple studies have now demonstrated that parental exposure to one pathogen can protect offspring against subsequent exposure to the same pathogen^1,13,19,20^. Several studies have also shown that parents exposed to a particular stress can produce progeny that are protected against the same stress, and that abiotic stress can provide transgenerational resistance to pathogenic infection^5,6,43,44^. Critically, it is not yet clear how the environment of the parent determines which stresses the offspring are protected against. Here, we show that inherited immunity to microsporidia can be activated by microsporidia and viral infection, as well as heavy metal stress, which all elicit a similar transcriptional response in the parents. Inherited immunity can also be activated through maternal somatic depletion of negative regulators of the transcriptional response. Indeed, activation of the transcriptional response within just a single generation in the intestine is sufficient to induce inherited immunity in offspring. Taken together, we have demonstrated that the signal for initiating inherited immunity against microsporidia is somatic induction of a shared maternal transcriptional stress response.

Inherited immunity phenotypes are often costly and can result in primed animals being more sensitive to other stresses^1,45^. Populations of *C. elegans* often develop within the same immediate environment, so parental infection is a good indicator of progeny exposure to microsporidia^46^. Here we find that progeny from microsporidia infected parents are smaller, carry fewer embryos, and are more sensitive to osmotic and heavy metal stress. While inherited immunity in response to microsporidia infection lasts only a single generation, it can be induced in every generation, and this may be a strategy to limit fitness trade-offs.

Immunity to microsporidia in *C. elegans* is thought to function through activation of the IPR and associated pathogen clearance^33,36^. Upregulation of the IPR results in immunity to both microsporidia and virus, but not bacteria^32^. Here, inherited immunity protects against microsporidia and bacteria, but not virus. Additionally, constant induction of the IPR genes in *pals-22* and *lin-35* mutants also greatly impacts animal development, whereas offspring with activated inherited immunity have less severe growth defects^32,47^. Our results suggest that although induction of the IPR is sufficient to activate inherited immunity, inherited immunity and IPR-mediated immunity are separate immune responses. We also show that inhibition of the RB ortholog *lin-35* provides immunity against microsporidia, both for the parents and their offspring. Although *lin-35* has been implicated in stress responses and negatively regulation of immune gene expression, this is the first example of *lin-35* mutants providing pathogen resistance^48,49^. RB is evolutionarily conserved, but in mammals acts as a positive regulator of antiviral immunity and immune cell development^50,51^.

Once the signal for inherited immunity is induced in the soma, the response must be transferred to developing progeny. Several inherited multigenerational responses that last for more than two generations are dependent on RNAi pathways^16,52^. Consistent with recent reports of responses that last only one or two generations being transmitted independent of these pathways, heritable immunity to microsporidia (which lasts a single generation) is not reliant on RNAi machinery^19,37^.

Being able to harness inherited immunity would provide a way to prevent infection of invertebrates^53^. Inherited immunity can be induced without the pathogen itself, by molecules that activate an immune response, and thus a similar approach could be used to combat microsporidia infections^54–56^. Inherited immunity could be employed to block infection of beneficial insects such as honey bees, and inhibition of this immunity could be used to improve the efficacy of microsporidia as biocontrol agents for locusts and other pests^57^. Although this is the first report of inherited immunity to microsporidia, mosquitoes whose parents were infected with microsporidia contained significantly fewer malaria parasites. This suggests that manipulation of these pathways could also be used to prevent invertebrate vectors of human disease from transmitting infection^58,59^.

## Materials and Methods

### 4.1. Worm maintenance

*C. elegans* strains were maintained at 21°C on nematode growth media (NGM) plates seeded with 10X OP50-1 *Escherichia coli*, as previously described (Brenner, 1974). Strains used in this study are listed in Supplemental Table 1. For maintenance and infection assays, 10X concentrates of OP50-1 were prepared by growing cultures to saturation in lysogeny broth (LB) at 37°C for 16-18 h. Populations were synchronized by washing worms off plates with M9 solution and bleaching with sodium hypochlorite / 1M NaOH until the embryos of gravid adults were released into solution. Eggs were washed three times with M9, resuspended in 5 ml M9, and rotated at 21°C for 18-24 h to allow embryos to hatch into L1s. For pelleting of live worms, animals were centrifuged in microcentrifuge tubes for 30 s at 1400xg.

**Supplemental Table 1.** *C. elegans* strains used in this study

| Strain name | Genotype | Source |
| --- | --- | --- |
| N2 | Wild-type, Bristol strain | Caenorhabditis Genetics Center (CGC) |
| JMC101 | *csr-1(tor67[csr-1 exon2::GFP::Flag IV:7958598])IV*<br>*csr-1(tor67[gfp::3xflag::csr-1]) IV* | (Ouyang et al., 2019) |
| DA465 | *eat-2(ad465) II* | (Raizen et al., 1995) |
| ERT054 | *jyls8[C17H1.6p::GFP, myo-2::mCherry] X* | (Bakowski et al., 2014) |
| ERT071 | *jyls14[F26F2.1p::GFP, myo-2::mCherry]* | (Bakowski et al., 2014) |
| ERT356 | *pals-22(jy1) III* | (Reddy et al., 2017) |
| WM27 | *rde-1(ne219) V* | CGC |
| MT10430 | *lin-35(n745) I* | CGC |
| ERT304 | *pals-22(jy1) III; jyls8[C17H1.6p::GFP, myo-2p::mCherry] X* | (Reddy et al., 2017) |
| AWR33 | *lin-35(n745) I; jyls8[C17H1.6p::GFP, myo-2::mCherry] X* | This study |
| ERT526 | *jySi37[pals-22p::pals-22::GFP, unc-119(+)] II; pals-22(jy1) unc-119(ed3) III* | (Reddy et al., 2017) |
| ERT527 | *jySi38[vha-6p::pals-22::GFP, unc-119(+)] II; pals-22(jy1) unc-119(ed3) III* | (Reddy et al., 2017) |
| ERT534 | *jySi39[unc-119p::pals-22::GFP, unc-119(+)] II; pals-22(jy1) unc-119(ed3) III* | (Reddy et al., 2017) |
| ERT539 | *jySi41[dpy-7p::pals-22::GFP, unc-119(+)] II; pals-22(jy1) unc-119(ed3) III* | (Reddy et al., 2017) |
| AWR35 | *lin-35(n745) I; unc-119(ed3) III; vrls55[ges-1p::lin-35::GFP::FLAG::lin-35 3'UTR]* | This study |
| AWR36 | *lin-35(n745) I; unc-119(ed3) III; vrls60[lin-35p::lin-35::GFP::FLAG::lin-35 3'UTR]* | This study |
| AWR37 | *lin-35(n745) I; unc-119(ed3) III; vrls93[mex-5p::lin-35::GFP::FLAG::lin-35 3'UTR]* | This study |
| AWR38 | *lin-35(n745) I; unc-119(ed3) III; vrls56[pie-1p::lin-35::GFP::FLAG::lin-35 3'UTR]* | This study |
| AWR45 | *pals-22(kea8[pals-22::GFP::degron]) III* | This study |
| AWR59 | *keaSi10[rpl-28p::TIR1::mRuby::unc-54 3'UTR +Cbr-unc-119(+)] II; pals-22(kea8[pals-22::GFP::degron]) III* | This study |
| AWR61 | *keaSi11[vit-2p::TIR1::mRuby::unc-54 3'UTR +Cbr-unc-119(+)] II; pals-22(kea8[pals-22::GFP::degron]) III* | This study |
| AWR64 | *kea15[rgef-1p::TIR1::mRuby::unc-54 3'UTR + Cbr-unc-119(+)] II; pals-22(kea8[pals-22::GFP::degron]) III* | This study |
| AWR62 | *keaSi9[myo-3p::TIR1::mRuby::unc-54 3'UTR +Cbr-unc-119(+)] II; pals-22(kea8[pals-22::GFP::degron]) III* | This study |
| AWR63 | *keaSi12[dpy-7p::TIR1::mRuby::unc-54 3'UTR + Cbr-unc-119(+)] II; pals-22(kea8[pals-22::GFP::degron]) III* | This study |
| AWR41 | *lin-35(kea7[lin-35p::degron::GFP::lin-35]) I* | This study |
| AWR54 | *lin-35(kea7[lin-35p::degron::GFP::lin-35]) I; ieSi57[eft-3p::TIR1::mRuby::unc-54 3'UTR + Cbr-unc-119(+)] II* | This study |
| AWR56 | *lin-35(kea7[lin-35p::degron::GFP::lin-35]) I; ieSi64[gld-1p::TIR1::mRuby::gld-1 3'UTR + Cbr-unc-119(+)] II* | This study |
| YY538 | *hrde-1(tm1200) III* | CGC |
| WM156 | *R04A9.2(tm1116) X* | CGC |
| YY1083 | *wago-4(tm1019) II* | (Xu et al. 2018) |
| WM160 | *sago-1(tm1195) V* | CGC |
| WM154 | *sago-2(tm894) I* | CGC |
| Unnamed | *ppw-1 (tm914) I* | (The C. elegans Deletion Mutant Consortium et al., 2012) |
| NL2098 | *rrf-1(pk1417) I* | CGC |
| MT17463 | *set-25(n5021) III* | CGC |
| AU0066 | *pmk-1(km25) IV* | CGC |

### 4.2. Construction of transgenic strains

For strains with tissue-specific expression of *lin-35* in a *lin-35* mutant background, MT10430 was crossed to DP38. The resulting *lin-35(n745)* I; *unc-119(ed3)* III double mutant was then crossed to YL398, YL402, YL409, and YL468^47^

SapTrap was used to construct a repair template for introducing the auxin inducible degron tag via CRISPR^60^. Briefly, the ∼500-600 bp regions immediately upstream of the *pals-22* start codon or downstream of the *lin-35* stop codon were PCR amplified from genomic N2 DNA to act as homology arms. These were cloned into the pDD379 backbone, along with sgRNA and other SapTrap-SEC kit plasmids (Addgene)^61^. The repair construct and co-injection markers were microinjected into N2 worms and hygromycin resistant rollers were selected. The SEC was then excised via heat shock^62^.

Strains carrying *vit-2p*::TIR1, *rgef-1p*::TIR1, *myo-3p*::TIR1, *dpy-7p*::TIR1, and *rpl-28p*::TIR1 were generated by MosSCI^63^. Repair constructs were generated using Multisite LR Gateway cloning using pCFJ150 as LR entry vector and microinjected into strain EG6699.

All strains generated for this study were outcrossed to N2 at least three times before use. Plasmids and primers used in this study are listed in Supplemental Table 2.

### 4.3. Preparation of microsporidia spores

*N. parisii* spores were prepared as described previously^26^. In brief, microsporidia spores were used to infect large populations of *C. elegans* N2 worms. Infected worms were then harvested, mechanically disrupted using 1 mm diameter Zirconia beads (BioSpec), and the lysate filtered through 5 μm filters (Millipore Sigma**™**) to remove nematode debris. Spores preparations were assayed for bacterial growth and those that were free of contaminating bacteria were stored at −80°C. Each assay was performed using spores of the same batch. In total, five batches were employed in this study.

### 4.4. Microsporidia infection assays

#### 4.4.1. Basic infection and priming assays

Synchronized populations of L1 worms were mixed with 1 ml of 10X OP50-1 alone as uninfected controls, or 1 ml of 10X OP50-1 supplemented with *N. parisii* spores, and plated on 10 cm NGM plates (doses defined in Supplemental Table 3 below). For priming assays, 2,500 P0 animals were infected with a ‘low dose’ of *N. parisii* such that >90% of animals were infected at 72 hpi and >80% of animals were fertile in order to generate a sufficient yield of primed F1 embryos for subsequent experiments (Figure S1). At 72 hpi, worms were collected and washed three times in 1 ml M9. To measure infection in the parental generation, 10% of P0 worms were fixed and parasite burden and gravidity assessed. The remaining 90% of worms were bleached to obtain naïve or primed embryos from uninfected or infected adults, respectively.

Following hatching in M9, 1,000 naïve or primed F1 animals were challenged at the L1 stage on a 6 cm plate with a high dose of *N. parisii*. This resulted in ∼10-20% of naïve animals being gravid, so that fitness increases or decreases would be easily detectable. At 72 hpi, worms were fixed, stained with DY96, and gravidity and infection assessed by microscopy.

**Supplemental Table 3.**
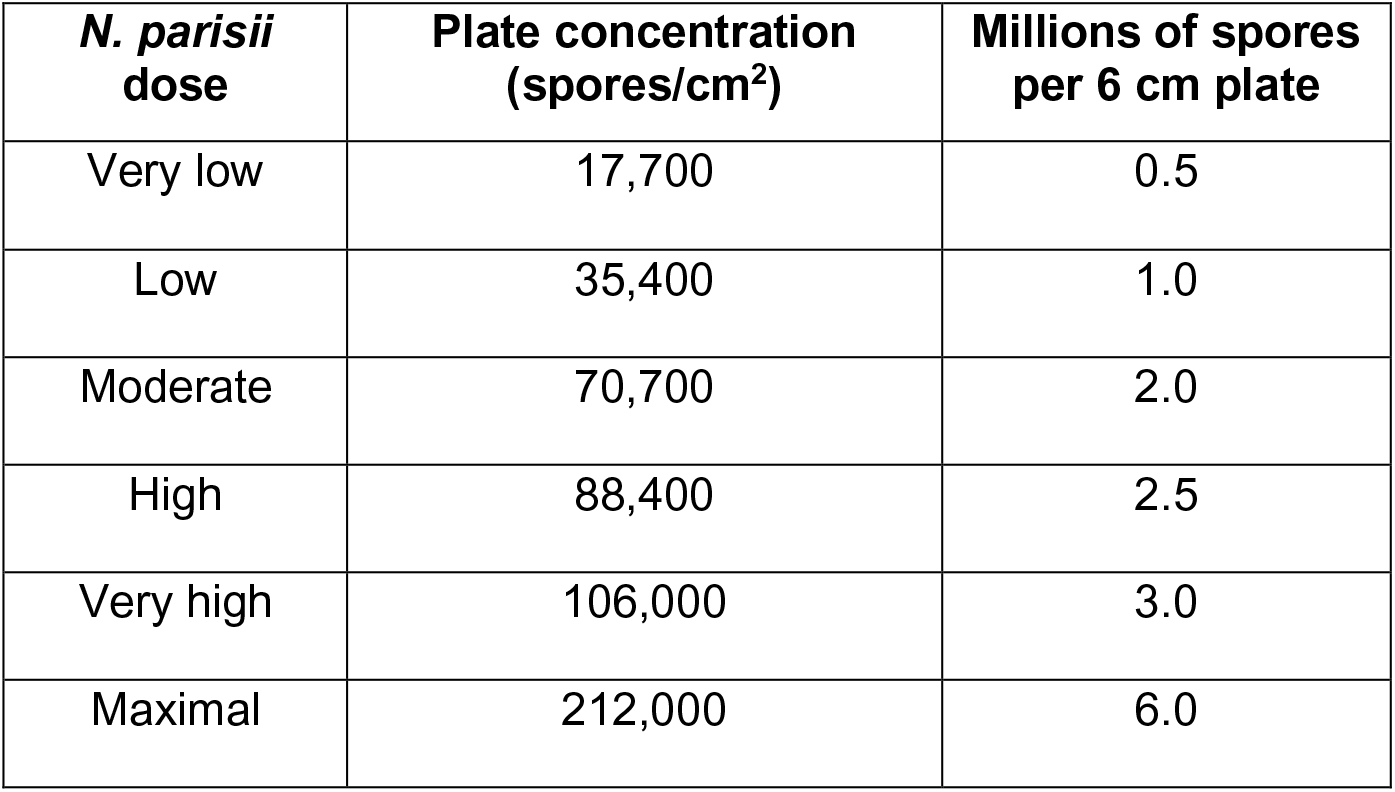
Doses of *N. parisii* used in this study

| ***N. parisii***<br>**dose** | **Plate concentration**<br>**(spores/cm<sup>2</sup>)** | **Millions of spores**<br>**per 6 cm plate** |
| --- | --- | --- |
| Very low | 17,700 | 0.5 |
| Low | 35,400 | 1.0 |
| Moderate | 70,700 | 2.0 |
| High | 88,400 | 2.5 |
| Very high | 106,000 | 3.0 |
| Maximal | 212,000 | 6.0 |

#### 4.4.2. Assessment of parental infection duration on inherited immunity

To test how long animals must be infected to transmit immunity to offspring, 2,500 P0 animals were either not infected or infected with a low dose of *N. parisii* at (i) the L1 stage, (ii) the L2/L3 stage after 24 h rest on 10X OP50-1 or (iii) the L4 stage after 48 h rest on 10X OP50-1. At 72 h post L1, P0 populations were bleached to release F1 embryos. Naïve and primed F1 animals were tested for inherited immunity as described in **Section 4.4.1**.

#### 4.4.3. Assessment of inherited immunity through development

To test inherited immunity throughout development, naïve or primed worms were obtained as described in **Section 4.4.1**. Next, 1,000 naïve or primed animals were plated on 10X OP50-1 supplemented with a maximal dose of *N. parisii* spores at (i) the L1 stage, (ii) the L2/L3 stage after 24 h rest on 10X OP50-1 or (iii) the L4 stage after 48 h rest on 10X OP50-1. At 30 mpi, animals were fixed, stained with MicroB FISH probe to detect *N. parisii* 18S RNA, and the number of sporoplasms quantified by microscopy.

#### 4.4.4. Assessment of spores in the gut and host cell invasion by *N. parisii*

To assay host cell invasion by microsporidia and the presence of spores in the gut, we conducted short infection assays ^24^. Naïve or primed F1 animals were obtained as described in **Section 4.4.1**. Next, 1,000 naïve or primed animals were mixed with 10X OP50-1 and a maximal dose of *N. parisii* spores at the L1 stage and plated on NGM. At 30 mpi, animals were fixed and stained with DY96 and MicroB FISH probe, and the number of spores or sporoplasms quantified by microscopy.

#### 4.4.5. Assessment of host clearance of *N. parisii*

To assay host clearance of microsporidia, we conducted pulse infection assays. For this, naïve or primed F1 animals were obtained as described in **Section 4.4.1**. Next, 2,000 naïve or primed animals were mixed with 10X OP50-1 and a very high dose of *N. parisii* spores at the L1 stage and plated on NGM. At 3 hpi, populations were split and half the animals fixed. The remaining animals were maintained in the absence of spores until a 24 h end point before fixing. Animals were stained with MicroB FISH probe, and the number of sporoplasms at each time point quantified by microscopy.

#### 4.4.6. Assessment of *N. parisii-*induced immunity over multiple generations

To assess immunity to microsporidia over multiple generations, we performed an ancestral infection assay (Figure S5C). Here, 2,500 animals were infected at the L1 stage with *N. parisii* for one, two or three successive generations (P0, F1, F2). Each infection period lasted 72 h before treating with sodium hypochlorite solution to obtain the next generation of embryos. F3 L1 populations were split and either subject to infection testing or maintained under non-infection conditions for the collection and subsequent testing of F4 offspring. To test immunity, F3 and F4 larvae were exposed to a high dose of *N. parisii* at the L1 stage. At 72 hpi, F3 and F4 animals were fixed and stained with DY96 to visualize *N. parisii* spores.

### 4.5. Heat-killed spore infection assay

Synchronized populations of 2,500 L1 worms were mixed with (i) 10X OP50-1 alone, or 10X OP50-1 supplemented with a low dose of either (ii) live or (iii) heat-killed *N. parisii* spores, and plated on NGM. For heat killing, live spores were treated at 65°C for 10 min. At 72 hpi, parental generations were bleached to obtain naïve embryos or embryos primed with heat-killed or live spores. Inherited immunity to *N. parisii* was tested as described in **Section 4.4.1**.

### 4.6. *P. aeruginosa* infection assays

Strains of wild-type or dsRed-expressing *P. aeruginosa* PA14 were used to assay survival and bacterial burden, respectively. Strains were grown in 3 ml LB overnight from a single bacterial colony. A volume of 20 μl of bacterial culture was used to seed 3.5 cm slow-killing plates for survival assays^35^. A volume of 50 μl of bacterial culture was used to seed 6 cm slow-killing plates for bacterial burden assays. To prevent *C. elegans* pathogen avoidance behaviour ^64^, bacterial culture was spread to ensure plates were fully covered with *P. aeruginosa*. Seeded plates were incubated for 24 h at 37°C for growth of bacterial lawns and maintained for a further 24 h at room temperature (RT) prior to infection assays. Naïve and primed worms were obtained as in **Section 4.4.1**. To assay survival, 20-30 naïve or primed L1 worms were transferred to 3.5 cm wild-type PA14-seeded plates and incubated at 25°C for 84 h. Worms that failed to respond to pressure from a metal pick were considered non-viable. To assay bacterial burden, 1,000 naïve or primed L1 worms were plated on 6 cm dsRed PA14-seeded plates, and incubated at 25°C for 48 h. At 48 hpi, worms were collected from plates, washed three times in M9, and fixed and bacterial burden assessed by microscopy.

### 4.7. Orsay virus infection assays

Orsay virus filtrate was prepared as described previously^29^. In brief, plates of Orsay virus infected animals were maintained until starvation. Virus shed by infected worms was collected by washing plates with M9, passing through 0.22 μm filters (Millipore Sigma**™**), and stored at −80°C. For infections to test resistance to Orsay virus, naïve and primed worms were obtained as in **Section 4.4.1**. Next, 1,000 naïve or primed L1s were mixed with 100 μl of 10X OP50-1 and 500 μl of the viral filtrate, then plated on 6 cm NGM. At 16 hpi, animals were fixed and FISH stained to assess infection status.

To generate Orsay-primed animals, 2,500 L1s were mixed with 1 ml of 10x OP50-1 and 500 μl of viral filtrate. At 72 hpi, animals were collected and bleached to obtain primed embryos. Inherited immunity to *N. parisii* was tested as described in **Section 4.4.1**.

### 4.8. Bead-feeding assay

Naïve or primed L1 worms were obtained as described in **Section 4.4.1**. Next,1,000 animals were placed on 6 cm plates in 400 μl total volume of M9 containing 10% (v/v) 10X OP50-1 and 4% (v/v) 0.2 μm green fluorescent polystyrene beads (Degradex Phosphorex). Where spores were included for the 3 h time point, a maximal dose of *N. parisii* spores was used. After 30 min or 3 h, animals were fixed, and bead ingestion assessed by microscopy.

### 4.9. Osmotic stress assays

Osmotic stress adaptation assays were performed as previously described^65^. Synchronized populations of 2,500 L1 worms were plated on either standard NGM (50 mM salt) plates together with (i) 10X OP50-1 alone or (ii) 10X OP50-1 supplemented with a low dose of *N. parisii*, or (iii) on NGM containing an elevated concentration of salt (250 mM) together with 10X OP50-1. At 80 hpi, parent populations were bleached to obtain naïve, infection-primed or salt-primed embryos. To test resistance to osmotic stress, 1,000 embryos were plated on NGM containing a high concentration of salt (420 mM). The percentage of embryos that had hatched by 48 h was quantified using light microscopy.

### 4.10. Cadmium assays

We first confirmed that cadmium exposure induced IPR gene expression in *C. elegans*. This was determined by plating ERT054 and ERT071 fluorescent reporter worm strains on 50 mM calcium chloride and observing increased GFP expression in exposed worms, relative to unexposed controls. Synchronized populations of 2,500 L4 worms were plated on either (i) standard NGM together with 10X OP50-1 alone or (ii) 10X OP50-1 supplemented with a low dose of *N. parisii*, or (iii) NGM containing 50 mM cadmium chloride together with 10X OP50-1. After 24 h, parent populations were bleached to obtain naïve, infection-primed or cadmium-primed embryos. Immunity to *N. parisii* was tested as described in **Section 4.4.1**. Immunity to *P. aeruginosa* was tested as described in **Section 4.6**. To test for protection against cadmium stress, 1,000 F1 larvae were maintained on standard NGM with 10X OP50-1 until L4 stage, then plated on NGM containing 50 mM cadmium chloride. After 24 h, animals were fixed and stained with DY96 for embryo counting.

### 4.11. Auxin-inducible depletion experiments

Auxin plates were prepared by adding auxin stock solution (400 mM auxin [Alfa Aesar] in ethanol) to NGM, for a final concentration of 200 uM auxin, immediately before pouring plates. Control plates were prepared by adding ethanol to NGM, for a final concentration of 0.15%. Auxin plates were stored in the dark at 4°C and used within one month.

Embryos obtained from bleaching gravid adults were plated on auxin or ethanol control plates following M9 washes. 10X OP50-1 was added to plates 18-24 h after plating to allow embryos to hatch and synchronize. Worms were bleached 72 h post L1 arrest and F1 immunity to *N. parisii* tested as described in **Section 4.4.1**.

### 4.12. Cross-progeny generation

For testing maternal effects, 50 L4 ERT054 males and 30 L4 hermaphrodites were plated on a 3.5 cm NGM plate for mating. To test paternal effects, L4 mutant males carrying the *jyIs8* transgene were crossed with L4 N2 hermaphrodites (Figure 5C). After 22-24 h, animals were bleached to obtain embryos and F1 immunity to *N. parisii* tested as described in **Section 4.4.1**. During quantification, the presence of *myo-2p::mCherry* was used to distinguish cross-progeny from self-progeny.

### 4.13. Fixation

For visualization of germlines of JMC101 (GFP::3xFLAG::CSR-1) animals, bead-feeding assays, and staining of *N. parisii* or Orsay virus with FISH probes, worms were fixed in 1 ml 4% paraformaldehyde (PFA) in phosphate buffered saline (PBS) containing 0.1% Tween-20 (PBST), for 20 min at RT or overnight at −20°C. For *P. aeruginosa* burden assays and DY96 staining, worms were fixed in 1 ml acetone for 10 min at RT, or overnight at 4°C. For pelleting of worms during fixing protocols, animals were centrifuged in microcentrifuge tubes for 30 s at 10,000xg

### 4.14. Staining with Direct Yellow 96 (DY96)

To assay parasite burden and worm gravidity, microsporidia spores and embryos were visualized with the chitin-binding dye DY96. Acetone-fixed animals were washed twice in 1 ml PBST, resuspended in 500 μl staining solution (PBST, 0.1% sodium dodecyl sulfate [SDS], 20 ug/ml DY96), and rotated at 21°C for 30 min in the dark. Stained worms were resuspended in 20 μl EverBrite™ Mounting Medium (Biotium) and mounted on slides for imaging. For pelleting of worms during staining protocols, animals were centrifuged in microcentrifuge tubes for 30 s at 10,000xg.

### 4.15. Fluorescence in Situ Hybridization (FISH) assays

For FISH staining of *N. parisii* 18S rRNA or Orsay virus RNA, worms were fixed in PFA as above and washed twice in 1 ml PBST. Worms were then washed once in 1 ml hybridization buffer (900 mM NaCl, 20 mM Tris pH 8.0, 0.01% SDS), and incubated overnight at 46°C in 100 μl hybridization buffer containing 5-10 ng/μl of FISH probe conjugated to a Cal Fluor 610 dye (LGC Biosearch Technologies). 5 ng/μl MicroB (ctctcggcactccttcctg)^27^ was used to detect *N. parisii* 18S RNA. 10 ng/μl of Orsay 1 (gacatatgtgatgccgagac) and Orsay 2 (gtagtgtcattgtaggcagc) mixed 50:50 was used to detect Orsay virus. Stained animals were washed once in 1 ml wash buffer (hybridization buffer containing 5 mM EDTA) and incubated in 500 μl fresh wash buffer for a further 30 min at 46°C. Worms were resuspended in 20 μl EverBrite™ Mounting Medium (Biotium) and mounted on slides for imaging. To pellet worms during staining protocols, animals were centrifuged in microcentrifuge tubes for 30 s at 10,000xg. For co-staining of *N. parisii* RNA with DY96, wash buffer was supplemented with 20 ug/ml DY96 for the final 30 min incubation.

### 4.16. Microscopy & image analysis

For measurement of worm body size, quantification of embryos and gravid or infected animals, as well as *N. parisii* or *P. aeruginosa* burden, worms were imaged using an Axioimager 2 (Zeiss). Worms carrying 1 or more embryos were considered gravid, worms possessing any quantity of DY96 stained microsporidia spores were considered infected. Bead ingestion and precise pathogen burdens were determined using ImageJ/FIJI^66^; here, each worm was defined as an individual ‘region of interest’ and fluorescence from GFP (DY96-stained microsporidia, or GFP beads) or dsRed (dsRed-expressing PA14) subject to ‘threshold’ and ‘measure area percentage’ functions on ImageJ. For DY96-stained samples. images were thresholded to capture the brighter signal from microsporidia spores, while eliminating the dimmer GFP signal from worm embryos. Final values are given as % fluorescence for single animals.

### 4.17. Transcriptional analyses

WormExp^67^ was used to search for published expression data sets that have a significant overlap with the set of genes enriched in *N. parisii* infected animals.

FPKM values from RNA-Seq data were obtained for *pals-22*^32^ mutants and five *N. parisii* infection time points^36^ and fold changes were calculated. Fold-change values were obtained for *lin-35* mutant microarray data^68^ and replicates averaged. Log_2_ fold-changes greater than 2 or less than −2 were used to determine differentially expressed genes. Genes that were either up or down regulated in *lin-35*, *pals-22* and at least one infection time point were plotted as a heatmap with dendrograms using heatmap.2 function from gplots package in R with all arguments^69^ set to default except for trace which was set to none.

### 4.18. Statistical analyses

Unless otherwise stated, p-values were determined by 2-tailed unpaired Student’s t-test. All p-values not meeting significance requirements are displayed on Figures for clarity. These p-values were calculated using Prism software (GraphPad Software Inc.). Statistical significance is defined as p-value < 0.05, unless otherwise stated (i.e. when using Bonferroni correction for multiple testing).

## Acknowledgements

We thank Emily R. Troemel for her mentorship and support in allowing A.W.R to start this project as a postdoctoral fellow in her lab. We thank Malina A. Bakowski and Lianne B. Cohen for sharing initial results on the shared transcriptional similarity between *lin-35* mutants and microsporidia infection. We thank Andy P. Ryan and Kirthi C. Reddy for sharing initial findings on the shared transcriptional similarity between cadmium exposure and microsporidia infection. We thank Nicholas O. Burton for advice on the osmotic stress assay. We are grateful to Calvin Mok and Nicholas O. Burton for providing helpful comments on the manuscript. Additional *C. elegans* strains were provided by the *Caenorhabditis* Genetics Center, which is funded by the National Institutes of Health (NIH) Office of Research Infrastructure Programs Grant P40 OD010440. Schematics were created using BioRender.com. This work was supported by the Natural Sciences and Engineering Research Council of Canada (Grant #522691522691).

## Supplementary figures

**Figure S1.**
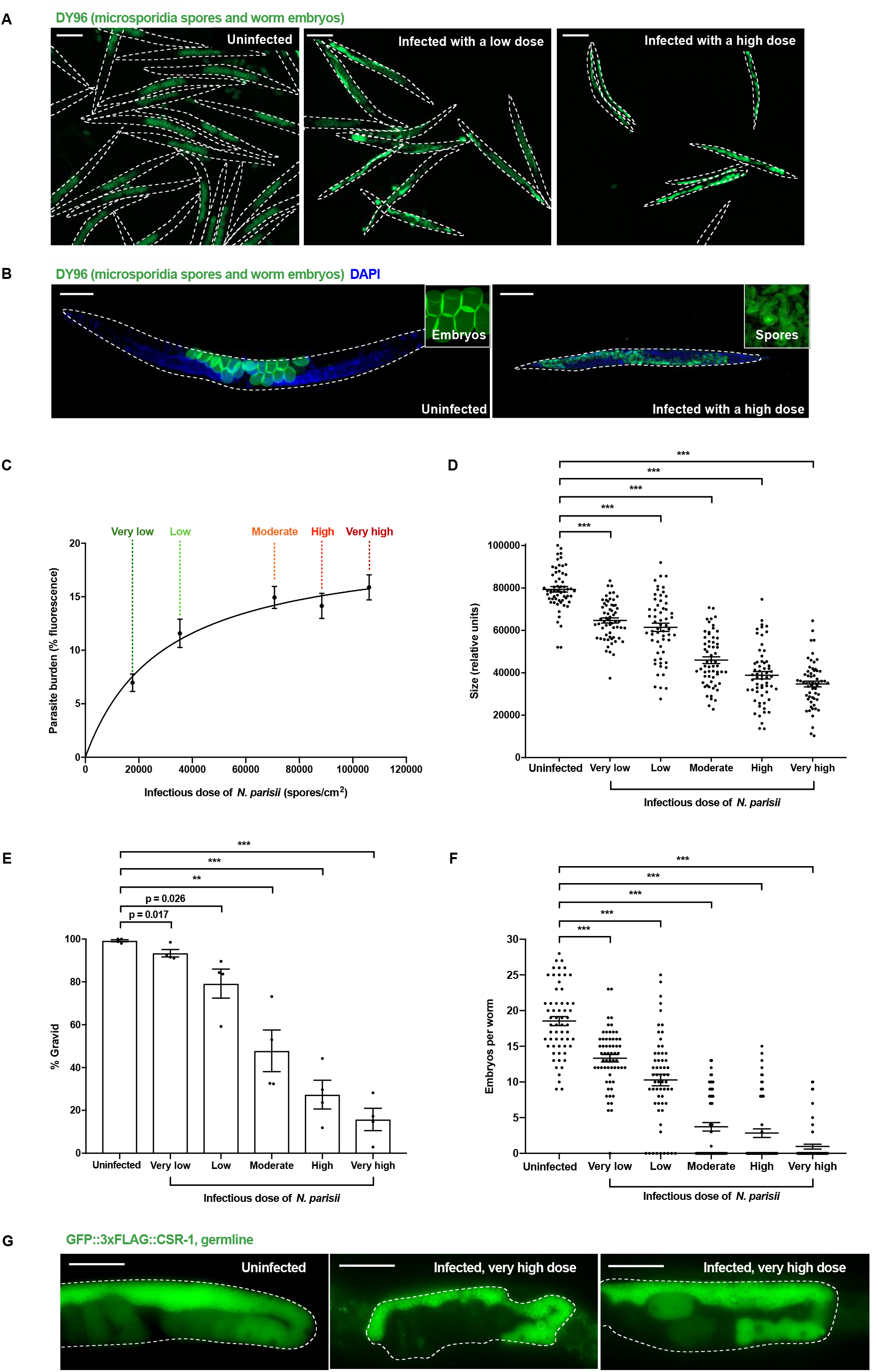
Infection of *C. elegans* by *N. parisii* severely impacts worm fitness. N2 *C. elegans* were uninfected or exposed to varying concentrations of *N. parisii* spores at the L1 stage (doses defined in methods). (A-F) At 72 hpi, animals were fixed and stained with DY96 to visualize both *N. parisii* spores and worm embryos. (A) Representative images of worm populations stained with DY96. Scale bars, 200 μm. (B) Representative images of individual worms stained with DY96 (green) and DAPI (blue). Left inset image shows DY96-stained embryos. Right inset image shows DY96 stained spores. Scale bars, 100 μm. (C) Images of DY96 stained worms were analysed and fluorescence from *N. parisii* spores thresholded to determine parasite burdens of individual worms (% of body filled with spores). Mean ± SEM (horizontal bars) is shown. Curve generated using the hyperbola nonlinear fit line function on GraphPad Prism software. Data pooled from 3 independent experiments using n = 20 worms per condition per experiment. (D) Images of worms were analysed, and the area of individual worms calculated. Each circle represents a measurement from a single worm. Mean ± SEM (horizontal bars) is shown. Data pooled from 3 independent experiments using n = 20 worms per condition per experiment. (E) Images of DY96 stained worms were analysed and worms possessing 1 or more embryos were considered gravid. Mean ± SEM (horizontal bars) is shown. Data pooled from 4 independent experiments using n = 35-195 worms per condition per experiment. (F) Images of DY96 stained worms were analysed and embryos per worm quantified. Each circle represents a count from a single worm. Mean ± SEM (horizontal bars) is shown. Data pooled from 3 independent experiments using n = 20 worms per condition per experiment. The p-values were determined by unpaired two-tailed Student’s t-test. (D-F) Significance with Bonferroni correction was defined as p < 0.01. **, p < 0.002; ***, p < 0.0002. (G) Transgenic *C. elegans* expressing a fluorescent germline protein (GFP::3xFLAG::CSR-1) were infected with *N. parisii* as above. Representative images of the germlines of individual worms. Scale bars, 50 μm.

**Figure S2.**
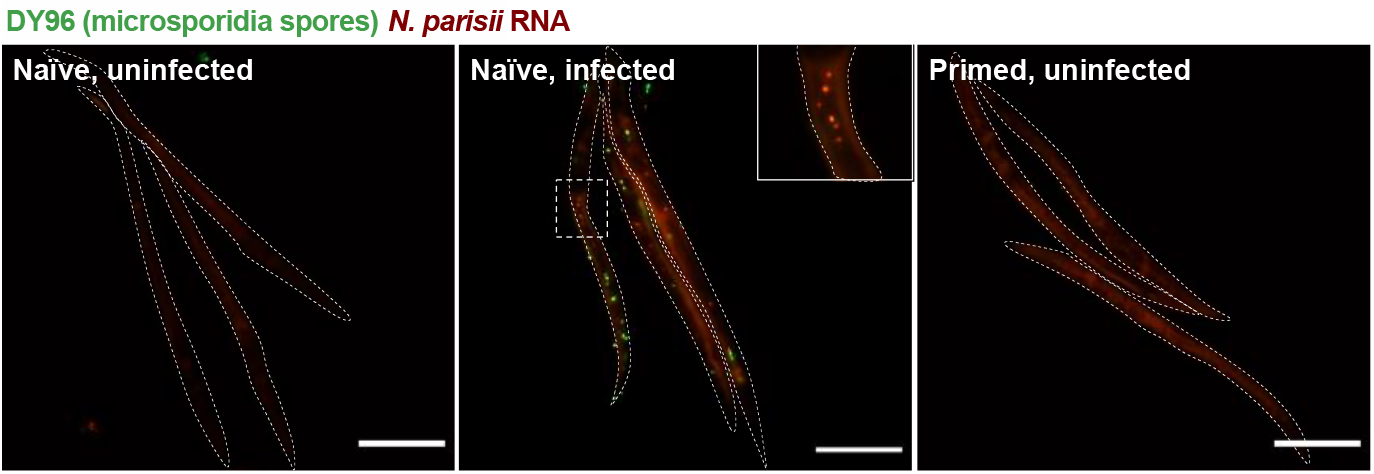
*N. parisii* infection of *C. elegans* is not vertically transmitted. P0 populations of N2 *C. elegans* were either not infected or infected with a moderate dose of *N. parisii* spores at the L1 stage (doses defined in methods). At 72 hpi, animals were treated with sodium hypochlorite solution to release F1 embryos. Naïve uninfected larvae served as a negative control for *N. parisii* detection. Naïve infected larvae exposed to a maximal dose of *N. parisii* served as a positive control for *N. parisii* detection. At 2 hpi F1 animals were fixed and stained with DY96 to detect *N. parisii* spores (green) as well as a FISH probe to detect *N. parisii* 18S RNA (red). *N. parisii* was never observed in primed, uninfected animals. Representative images of F1 populations are shown. Scale bars, 50 μm.

**Figure S3.**
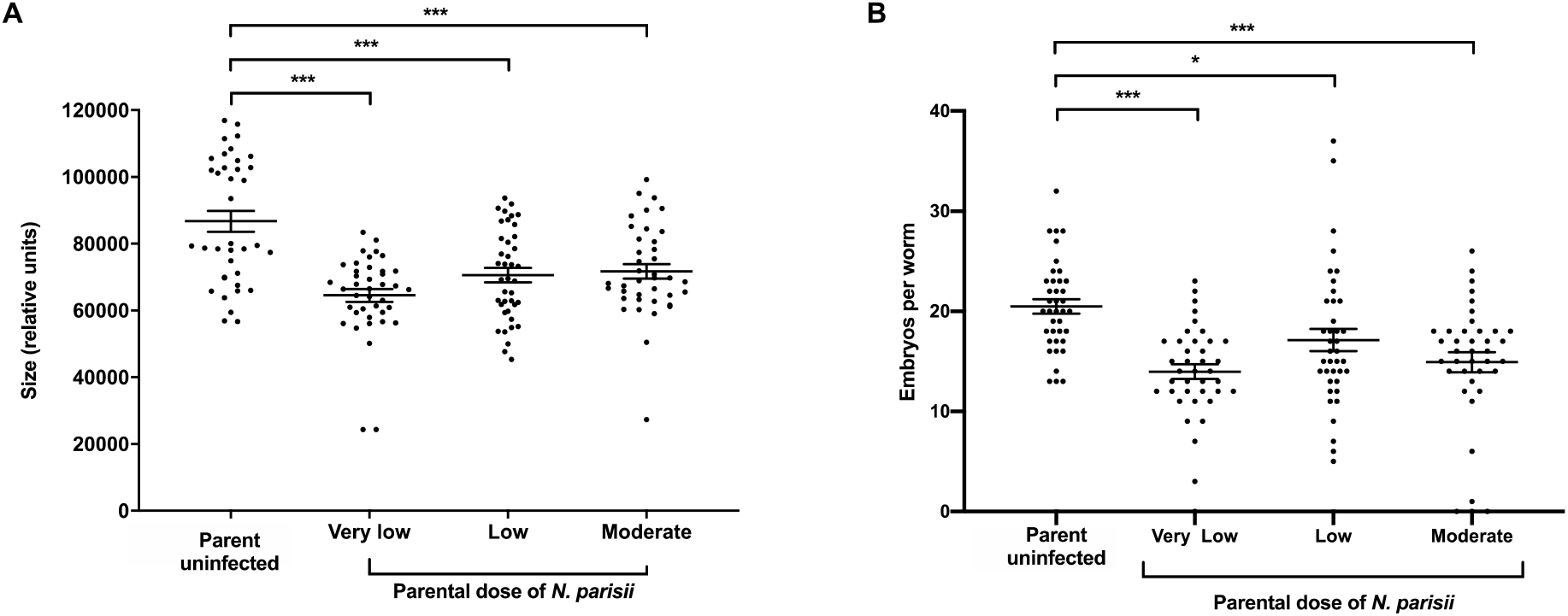
Progeny of *N. parisii* infected *C. elegans* are less fit when uninfected. P0 populations of N2 *C. elegans* were uninfected or exposed to varying concentrations of *N. parisii* spores at the L1 stage (doses defined in methods). At 72 hpi, animals were treated with sodium hypochlorite solution to release F1 embryos. Naïve or primed F1 L1 larvae were grown under non-infection conditions for 72 h then fixed and stained with DY96 to visualize worm embryos. (A) Images of F1 worms were analysed, and the area of individual worms calculated. Each circle represents a measurement from a single worm. Mean ± SEM (horizontal bars) is shown. Data pooled from 2 independent experiments using n = 17-20 worms per condition per experiment. (B) Images of DY96 stained F1 worms were analysed and embryos per worm quantified. Each circle represents a count from a single worm. Mean ± SEM (horizontal bars) is shown. Data pooled from 2 independent experiments using n = 20 worms per condition per experiment. The p-values were determined by unpaired two-tailed Student’s t-test. Significance with Bonferroni correction was defined as: *, p < 0.0166; **, p < 0.0033; ***, p < 0.00033.

**Figure S4.**
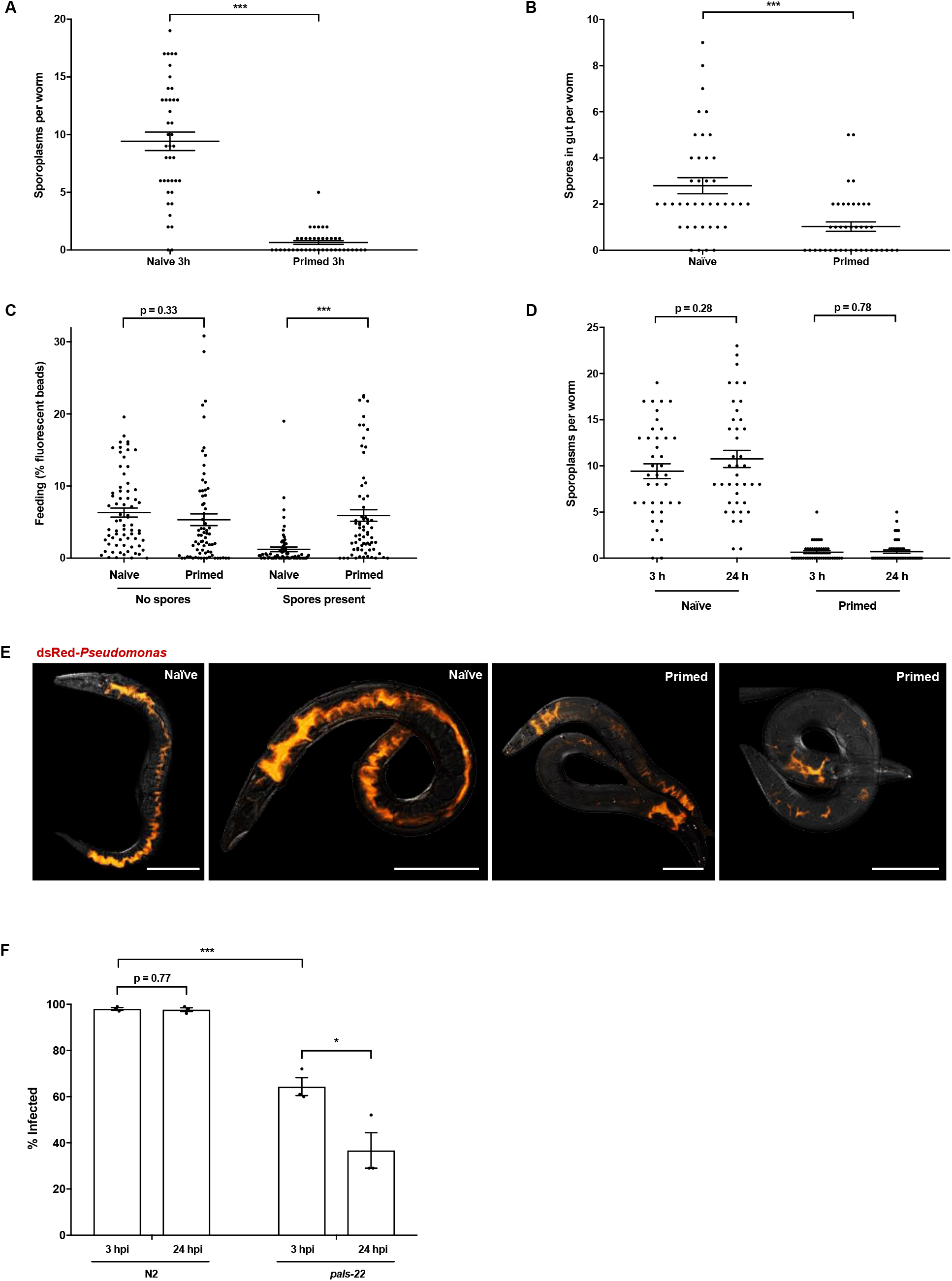
Inherited immunity protects primed larvae from infection. Inherited immunity reduces microsporidia and *Pseudomonas* in the intestine, whereas *pals-22 mutants* both reduce microsporidia invasion and cause microsporidia clearance. P0 populations of N2 *C. elegans* were either not infected or infected with a low dose of *N. parisii* spores at the L1 stage (doses defined in methods). At 72 hpi, animals were treated with sodium hypochlorite solution to release F1 embryos. (A-B) Naïve and primed F1 larvae were exposed to a very high dose of *N. parisii* at the L1 stage. At 3 hpi, animals were fixed and stained with DY96 to detect *N. parisii* spores (green) as well as a FISH probe to detect *N. parisii* 18S RNA and reveal intracellular sporoplasms (red). (A) The number of sporoplasms per animal was quantified by microscopy. Each circle represents a count from a single worm. Mean ± SEM (horizontal bars) is shown. Data pooled from 2 independent experiments using n = 20-26 worms per condition per experiment. (B) The number of spores per animal was quantified by microscopy. Each circle represents a count from a single worm. Mean ± SEM (horizontal bars) is shown. Data pooled from 2 independent experiments using n = 20 worms per condition per experiment. (C) Naïve and primed F1 larvae were fed on fluorescent beads at the L1 stage. After 3 hours, animals were fixed and imaged. Fluorescence from beads was thresholded to determine the amount of beads eaten by each worm (% of body filled with beads). Each circle represents a measurement from a single worm. Mean ± SEM (horizontal bars) is shown. Data pooled from 2 independent experiments using n = 20-30 worms per condition per experiment. (D) Naïve and primed F1 larvae were exposed to a very high dose of *N. parisii* at the L1 stage. At 3 hpi, populations were split and half of the animals fixed. The remaining animals were maintained in the absence of spores until a 24 h end point before fixing. Animals were stained with a FISH probe to detect *N. parisii* 18S RNA (red) and the number of sporoplasms per animal was quantified by microscopy. Each circle represents a count from a single worm. Mean ± SEM (horizontal bars) is shown. Data pooled from 2 independent experiments using n = 20 worms per condition per experiment. (E) Naïve and *N. parisii*-primed F1 L1 larvae were maintained on slow-killing plates with dsRed-*P. aeruginosa* and fixed at 48 hpi. Representative images of worms grown on dsRed-*P. aeruginosa* are shown. Scale bars, 100 μm. (F) N2 and *pals-22* F1 larvae were exposed to a very high dose of *N. parisii* at the L1 stage. At 3 hpi, populations were split and half of the animals fixed. The remaining animals were maintained in the absence of spores until a 24 h end point before fixing. Animals were stained with a FISH probe to detect *N. parisii* 18S RNA (red) and the number of sporoplasms per animal was quantified by microscopy. Three independent experiments with n = 100 worms per experiment. The p-values were determined by unpaired two-tailed Student’s t-test. Significance with Bonferroni correction was defined as p < 0.05. ***, p < 0.001.

**Figure S5.**
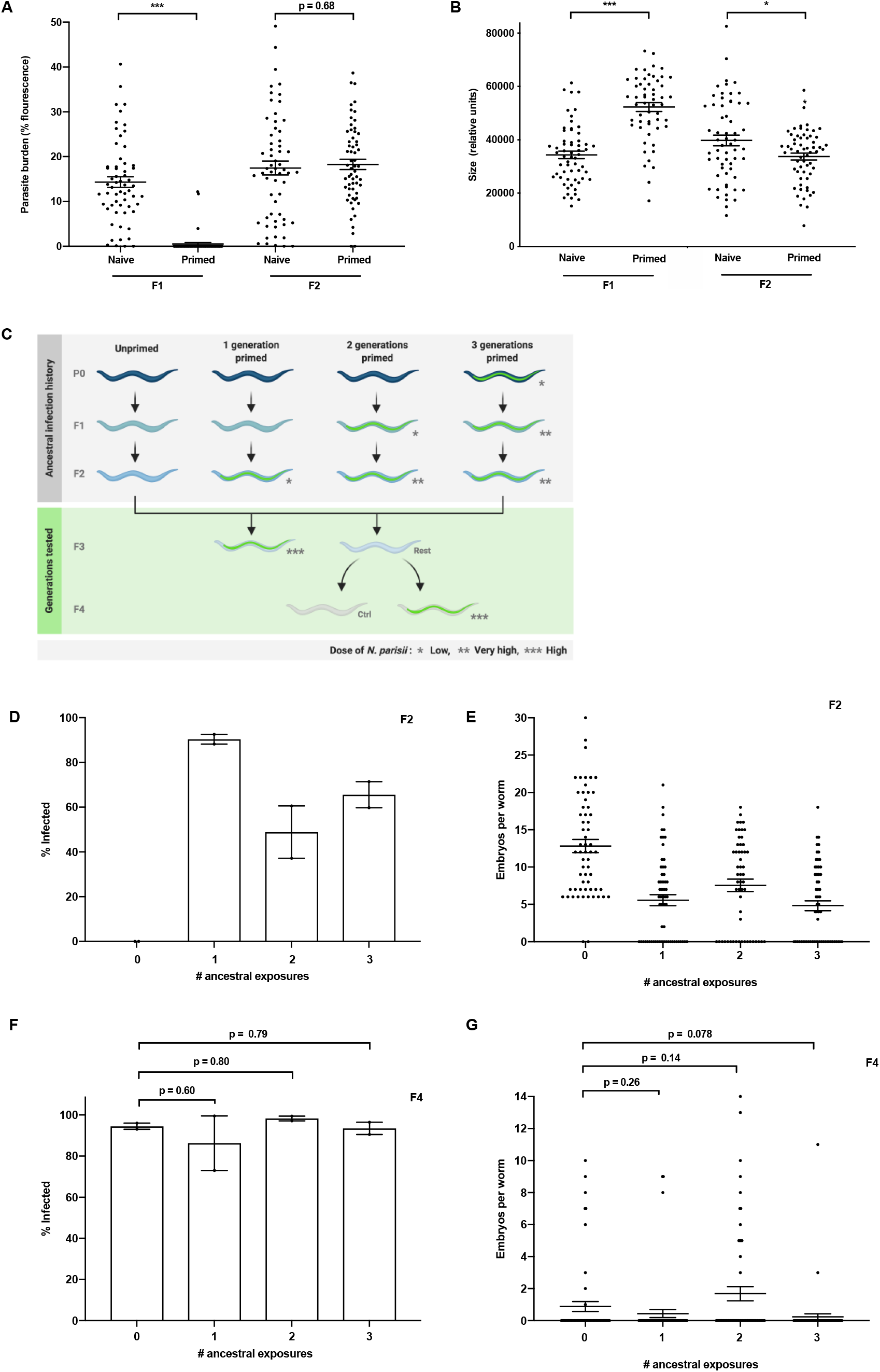
Inherited immunity in *N. parisii* primed *C. elegans* lasts a single generation. (A-B) P0 populations of N2 *C. elegans* were either not infected or infected with a low dose of *N. parisii* spores at the L1 stage (doses defined in methods). At 72 hpi, animals were treated with sodium hypochlorite solution to release F1 embryos. F1 L1 populations were then split and either subject to infection testing, or maintained under non-infection conditions for the collection and subsequent testing of F2 embryos. Both naïve and primed F1 and F2 larvae were exposed to a high dose of *N. parisii* at the L1 stage. At 72 hpi, F1 and F2 animals were fixed and stained with DY96 to visualize *N. parisii* spores. (A) Images of DY96 stained worms were analysed and fluorescence from *N. parisii* spores thresholded to determine parasite burdens of individual worms (% of body filled with spores). Each circle represents a measurement from a single worm. Mean ± SEM (horizontal bars) is shown. Data pooled from 3 independent experiments using n = 15-20 worms per condition per experiment. (B) Images of worms were analysed, and the area of individual worms calculated. Each circle represents a measurement from a single worm. Mean ± SEM (horizontal bars) is shown. Data pooled from 3 independent experiments using n = 15-20 worms per condition per experiment. (C) Schematic of ancestral infection history assay. N2 *C. elegans* (P0, F1, F2) were infected at the L1 stage with *N. parisii* for one, two or three successive generations. Each infection period lasted 72 h before treating with sodium hypochlorite solution to obtain the next generation of embryos. F3 L1 populations were split and either tested for immunity or maintained under non-infection conditions for the collection and subsequent testing of F4 offspring. For infection testing, F3 and F4 larvae were exposed to a high dose of *N. parisii* at the L1 stage. At 72 hpi, F3 and F4 animals were fixed and stained with DY96 to visualize *N. parisii* spores. (D) As the offspring of infected parents are resistant to infection, second (F1) or third (F2) generation doses were increased to ensure these primed animals still became infected. To determine infection status of F2 populations prior to testing of next generations, individual DY96 stained worms were imaged to determine infection status. Mean ± SEM (horizontal bars) is shown. Data pooled from 2 independent experiments using n = 35-133 worms per condition per experiment. (E) Images of DY96 stained F2 worms were analysed and embryos per worm quantified. Each circle represents a count from a single worm. Mean ± SEM (horizontal bars) is shown. Data pooled from 2 independent experiments using n = 25-30 worms per condition per experiment. (F) Individual DY96 stained F4 worms were imaged to determine infection status. Mean ± SEM (horizontal bars) is shown. Data pooled from 2 independent experiments using n = 37-200 worms per condition per experiment. (G) Images of DY96 stained F4 worms were analysed and embryos per worm quantified. Each circle represents a count from a single worm. Mean ± SEM (horizontal bars) is shown. Data pooled from 2 independent experiments using n = 30 worms per condition per experiment. The p-values were determined by unpaired two-tailed Student’s t-test. (A-B) Significance was defined as: *, p < 0.05; **, p < 0.01; ***, p < 0.001. (F-G) Significance with Bonferroni correction was defined as p < 0.016.

**Figure S6.**
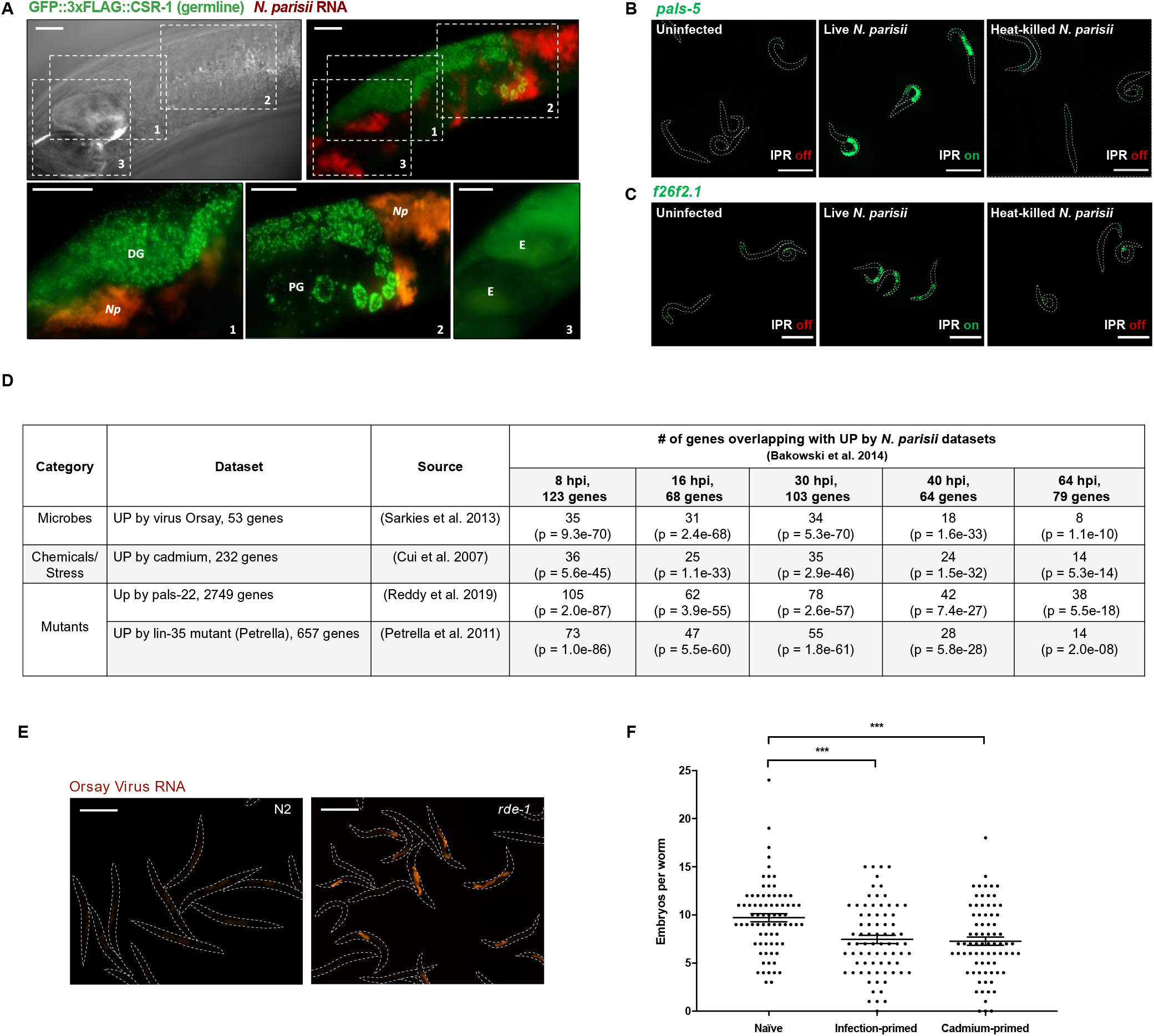
Transmission of inherited immunity requires the transcriptional response to microsporidia, which is mimicked by other environmental conditions and mutants. (A) Transgenic *C. elegans* expressing a fluorescent germline protein (GFP::3xFLAG::CSR-1) were infected with a low dose of *N. parisii* spores at the L1 stage (doses defined in methods). At 72 hpi, animals were fixed and stained with a FISH probe to detect *N. parisii* 18S RNA (red). Representative images of the germline of an infected worm are shown. Inset images: *Np*, *N. parisii*; (1) DG, distal gonad; (2) PG, posterior gonad; (3) E, embryo. Scale bars, 20 μm. (B-C) Transgenic *C. elegans* expressing GFP under IPR gene promoters were either uninfected or exposed to a low dose of heat-killed or live *N. parisii* spores at the L1 stage. At 5 hpi, animals were imaged. Scale bars, 100 μm. (B) Representative images of transgenic *pals-5p::gfp* worms. (C) Representative images of transgenic *f26f2.1p::gfp* worms. (D) Table comparing genes previously reported to be upregulated in *N. parisii* infected animals with gene expression changes in other published data sets. p-values calculated using Fisher’s Exact test. (E) N2 and *rde-1* mutant L1 larvae were infected with Orsay virus and fixed at 72 hpi. Representative images of worms stained with FISH probe to detect Orsay virus RNA. Scale bars, 400 μm. (F) P0 populations of N2 *C. elegans* were either untreated, exposed to 50 mM cadmium from the L4 stage, or infected with a low dose of *N. parisii* spores at the L4 stage. After 24 h, animals were treated with sodium hypochlorite solution to release F1 embryos. F1 larvae were exposed to 50 mM cadmium at the L4 stage. After 24 h, animals were fixed and stained with DY96 to visualize worm embryos. Images of DY96 stained worms were analysed and embryos per worm quantified. Each circle represents a count from a single worm. Mean ± SEM (horizontal bars) is shown. Data pooled from 3 independent experiments using n = 25-26 worms per condition per experiment. The p-values were determined by unpaired two-tailed Student’s t-test. Significance with Bonferroni correction was defined as p < 0.025. ***, p < 0.0005.

**Figure S7.**
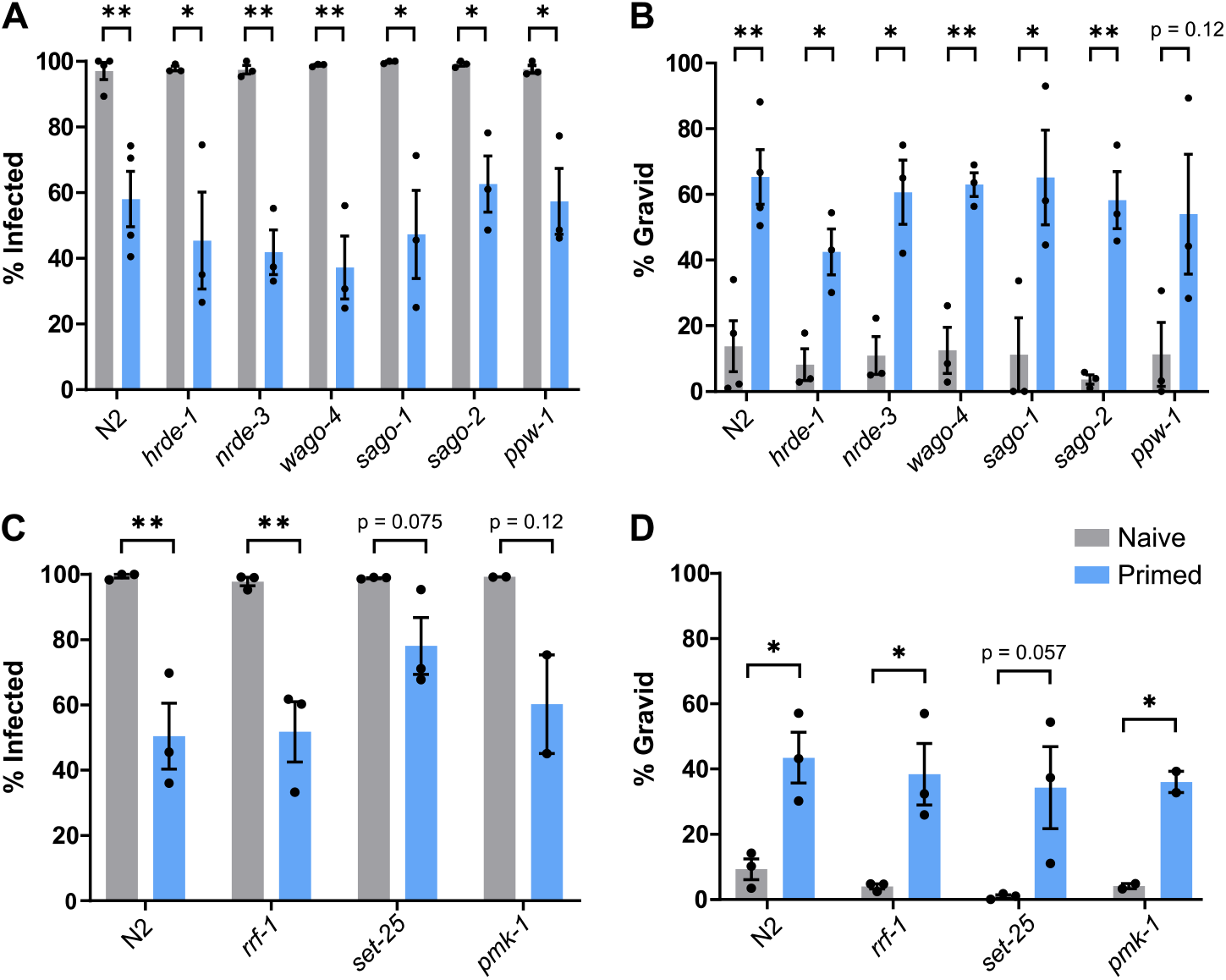
Intergenerational transmission of immunity does not depend on small-RNA inheritance factors, a histone methyltransferase, or the P38 MAP kinase pathway. (A-D) N2 or mutant P0 animals were either not infected or infected with a moderate dose of *N. parisii* spores at the L1 stage (doses defined in methods). At 72 hpi, animals were treated with sodium hypochlorite solution to release F1 embryos. Naïve or primed F1 L1 larvae were then infected with a high dose of *N. parisii*. Percentage of infected animals (A, C) and gravid animals (B, D) were quantified in the infected F1s at 72hpi. Mean ± SEM (horizontal bars) is shown. 2-5 independent experiments with n > 100 worms each. The p-values were determined by unpaired two-tailed Student’s t-test. Significance was defined as: *, p < 0.05; **, p < 0.01; ***, p < 0.001; ****, p < 0.0001

**Figure S8.**
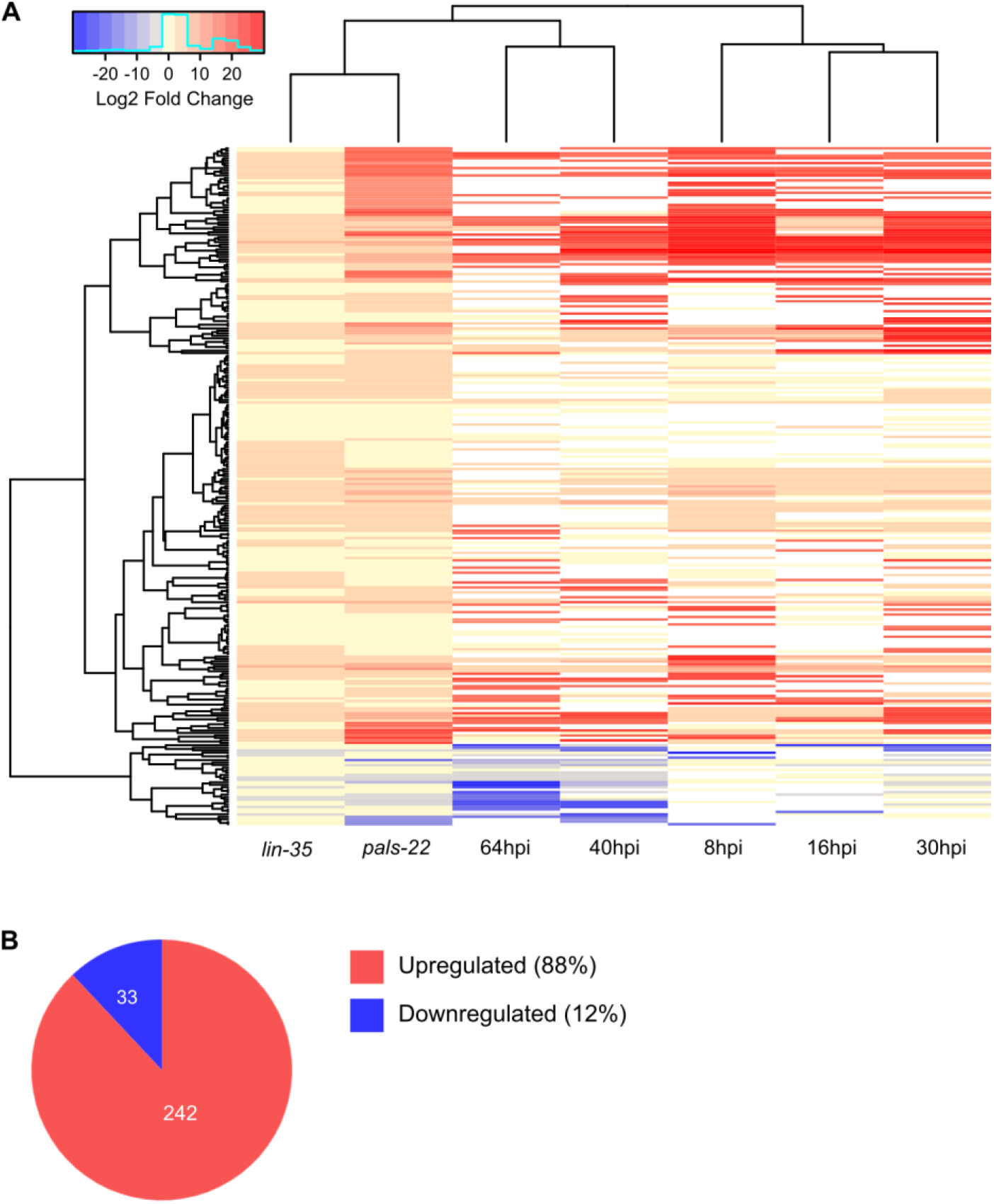
*N. parisii* infection induces many genes that are also upregulated in both *lin-35* and *pals-22* mutants. A shared transcriptional response was identified by determining genes that were either up or down regulated in both *lin-35* and *pals-22* mutants and at least one *N. parisii* infection time point. (A) Heat map showing cluster analysis of the shared transcriptional response with the fold change of each cell corresponding to scale at the top. White cells in heatmap represent gene not determined to be differentially expressed. (B) Fraction of shared genes that are either up or down regulated.

**Figure S9.**
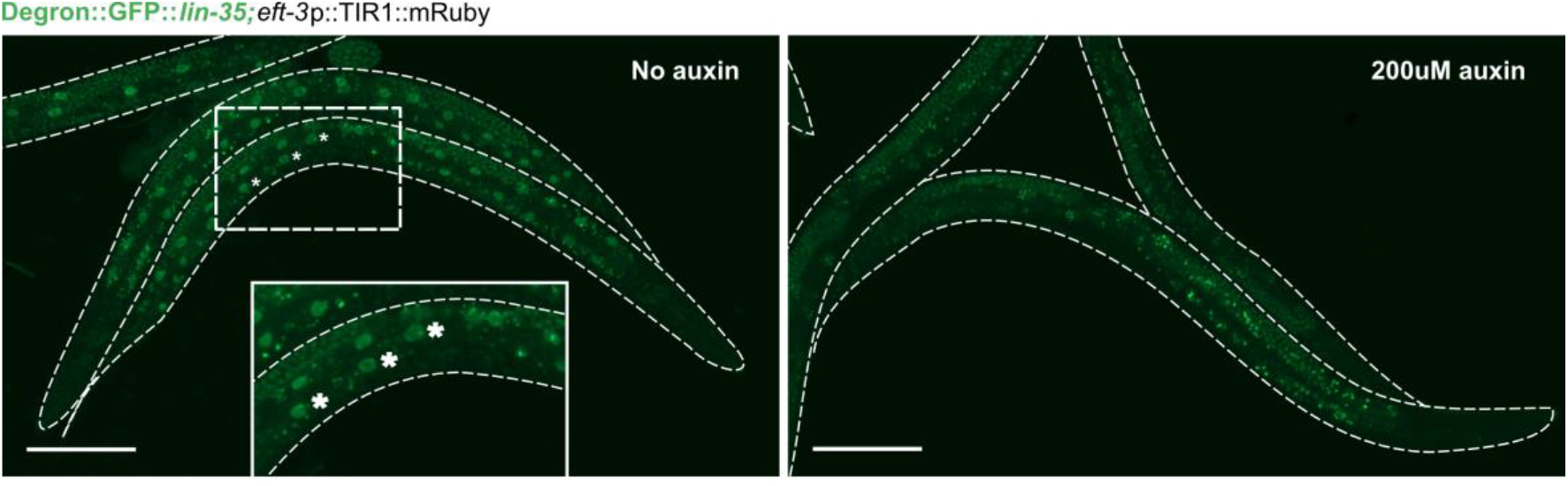
Degradation of *lin-35* in somatic tissues. Degron::GFP::*lin-35* worms with somatic TIR1 expression were grown on control plates, containing no auxin, or 200 uM auxin plates for degradation of LIN-35. Images of representative animals show GFP expression 48 h post-hatch on their respective plates. Asterisks mark intestinal nuclei GFP. Scale bars, 100 μm.

## Notes

### Competing Interest Statement

The authors have declared no competing interest.

